# Absolute voltage mapping using dynamic photocycle control

**DOI:** 10.64898/2026.07.17.739178

**Authors:** Daozheng Gong, Jinfan Zhang, Madeleine R. Howell, Xiang Wu, Bill Z. Jia, Adam E. Cohen

**Affiliations:** Department of Chemistry and Chemical Biology, Harvard University; Department of Physiology, University of California, San Francisco; Department of Physics, Harvard University

## Abstract

Fluorescent voltage indicators are widely used to report relative changes in membrane potential, but mapping absolute voltages remains difficult. Here we present Voltage Measurement by Activated Photocycles (VMAP), a simple method for absolute voltage imaging based on a photophysical switch between voltage-insensitive and sensitive indicator states. VMAP requires no specialized hardware or additional labeling, and is applicable across species, sample preparations, and microscope configurations. Using VMAP, we quantified drug-induced shifts in neuronal resting potential, revealed the emergence of bioelectric patterns during multi-day recordings of human iPSC populations, and created 3D membrane-potential maps across whole live zebrafish embryos. By making absolute voltage imaging accessible from cellular to organismal scales and from milliseconds to days, VMAP opens a route to mapping bioelectrical organization in complex living systems.

## Introduction

All cells maintain a membrane potential (V_m_), and voltage dynamics regulate biological processes across a vast range of scales, from millisecond action potentials to multicellular patterns that evolve during development, regeneration, and disease (*1–4*). Patch clamp recordings remain the gold standard for measuring membrane potential with high accuracy and temporal resolution. However, patch clamp is low throughput, lacks spatial resolution, and is incompatible with chronic recordings, making it difficult to measure membrane potential across complex tissues or *in vivo*.

Genetically encoded voltage indicators (GEVIs) enable optical recording of V_m_ dynamics from thousands of cells in parallel, and are thus becoming increasingly popular for reporting V_m_ dynamics (*5–8*). However, the voltage-dependent fluorescence, *F*_*v*_, of GEVIs is influenced by many confounding factors such as expression level, excitation intensity, cell morphology, and photobleaching (*9*). Normalizing fluorescence by a reference state (i.e. calculation of Δ*F*_*v*_/*F*_*0*_) can correct for variations in reporter density and illumination, but this quantity can only report absolute voltage if the membrane potential in the reference state, *F*_0_ is accurately known. This is rarely the case. As a result, voltage imaging is mostly limited to reporting relative voltage changes that occur faster than changes in any of the confounding variables (*10, 11*), and cannot, e.g. compare the resting potentials of adjacent cells.

A few approaches have been developed to measure absolute voltage optically, but each faces technical barriers. Fluorescence lifetime imaging microscopy (FLIM) quantifies the nanosecond-timescale fluorescence lifetime of GEVIs, but requires complex and specialized hardware and suffers from low throughput (*12–15*). Dual-label ratiometric imaging uses a second fluorescent marker as a reference baseline (*16–19*). This approach consumes much of the visible spectrum and is susceptible to artifacts caused by differential production or degradation of the two reporters. A simple, accessible method for absolute voltage imaging with uncompromised accuracy and broad compatibility could make bioelectrical mapping commonplace in biological research.

### Principle of voltage measurement by activated photocycle

The photocycles of some microbial rhodopsins contain V_m_-dependent kinetic steps that can support internally referenced measurements of absolute V_m_ (*20, 21*). Early applications, however, required multiple wavelengths to drive and probe the photocycle and were thus not practical for widespread use. Recently, we found that the voltage sensitivity of Voltron2 (*22*), a widely used FRET-opsin GEVI, arises from a photocycle intermediate, not the ground state (*23*). This photocycle feature reveals an intrinsic voltage-insensitive reference state and revives the possibility of using the opsin photocycle for absolute voltage imaging.

Here we introduce Voltage Measurement by Activated Photocycle (VMAP), a technically simple method for absolute voltage imaging. The principle of VMAP is based on the photocycle of indicators such as Voltron2 (Fig. 1A): in the dark, Voltron2 relaxes to a state whose fluorescence is insensitive to *V*_m_. The fluorescence of this state, *F*_i_, is probed by one or more sub-millisecond light pulses. A subsequent prolonged illumination step then drives the indicator into its voltage-sensitive manifold of states whose fluorescence *F*_*v*_ is modulated by V_m_. This interleaving of short and long illumination pulses is repeated for each absolute voltage measurement. The ratio *R*_v_ = *F*_*v*_/*F*_*i*_ cancels voltage-independent factors and provides a robust metric of absolute V_m_.

**Figure 1.**
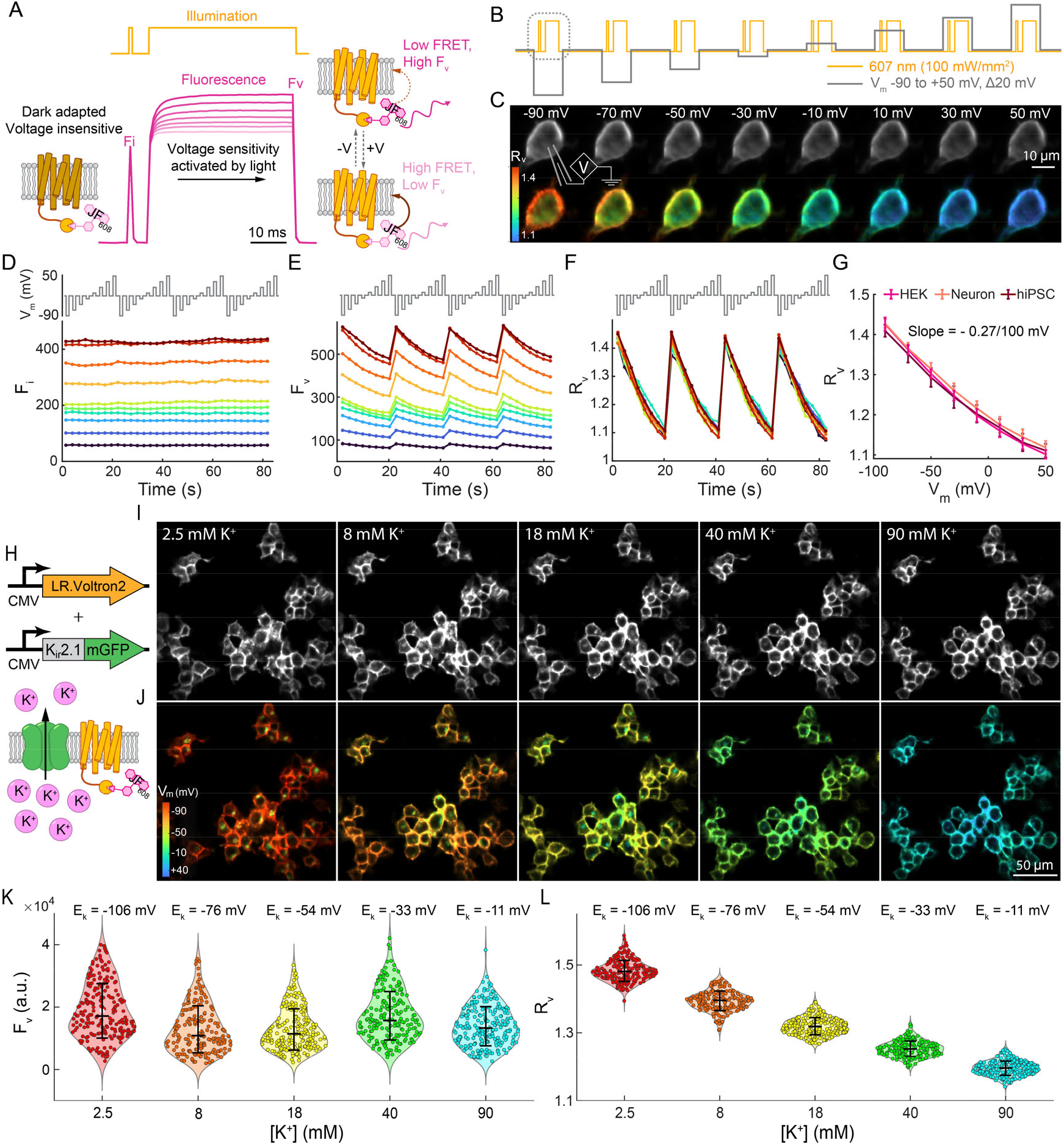
Absolute voltage imaging via VMAP. (A) Schematic of the VMAP principle, based on the Voltron2 photocycle. A brief illumination pulse probes the dark-adapted, voltage-insensitive fluorescence (*F*_i_). A longer illumination pulse populates the voltage-sensitive manifold, with fluorescence *F*_v_. (B) Voltage-clamp protocol with synchronized illumination used for VMAP measurements. The circled illumination pulses correspond to the protocol in (A). (C) Voltage-dependent VMAP images of a HEK cell expressing Voltron2-JF608. The ratio *R*_v_ = *F*_v_/*F*_i_ reports absolute membrane voltage. (D–F) Time courses of (D) *F*_i_, (E) *F*_v_, and (F) *R*_v_ during repeated voltage steps, measured in *n* = 10 HEK cells. (G) Calibration of *R*_v_ vs. membrane potential (*V*_m_) in HEK cells, neurons, and hiPSCs, showing an approximately linear relationship between −90 to −10 mV (slope ≈ -0.27 per 100 mV). Error bars represent mean ± s.d.. (H) Expression of K_ir_2.1 set the resting potential to the K^+^ Nernst potential. (I) Fluorescence images and (J) corresponding *R*_v_ maps at increasing extracellular [K^+^]. (K) Population distributions of raw fluorescence (F_v_) and (L) *R*_v_ across ∼200 cells per condition. Error bars represent mean ± s.d.. All data acquired on the spinning disk confocal microscope.

We optimized the VMAP protocol using Voltron2 expressed in HEK293 cells, labeled with the HaloTag ligand dye JF608, and imaged in a widefield epifluorescence microscope with illumination at 594 nm, 160 mW/mm^2^ (**Methods**, Table S1). We selected the JF608 dye for its brightness, photostability, bioavailability *in vivo* and red spectrum (permitting combination with optogenetic stimulation or imaging of blue/green reporters). We systematically varied the timing of the short and long pulses and the intervening dark intervals. The initial dark interval must be long enough to ensure relaxation of any previously populated voltage-sensitive states. The subsequent pulse for probing *F*_*i*_ must be short enough to avoid populating the voltage-sensitive state during the *F*_*i*_ measurement. A 150 ms dark interval followed by a 0.5 ms probe pulse yielded a residual voltage sensitivity *ΔF*_i_/*F*_i_ per 100 mV ≈ 1.3%, sufficiently small compared to the voltage-sensitive state (Δ*F*_*v*_/*F*_*v*_ per 100 mV ≈ 23%) to enable reliable normalization while still providing adequate signal in *F*_i_ (Fig. S1).

Upon onset of steady-state illumination, voltage sensitivity arose with a time constant of 4 ms. We typically measured *F*_*v*_ after at least 30 ms of steady-state illumination to ensure that illumination-dependent transients had fully relaxed. The long illumination step can be extended for quasi-continuous recordings of fast voltage dynamics, taking advantage of the sub-millisecond voltage response of Voltron2. In this mode, brief dark intervals and short pulses are interspersed at user-selected intervals to recalibrate *F*_*i*_. In some cases, the *F*_*i*_ measurement was contaminated by shot noise, due to the short (0.5 ms) exposure time. In these cases, we made a series of *F*_*i*_ measurements spaced by 100 ms. These were then averaged together before calculating *R*_v_ (Fig. S1B). This procedure mitigated the effect of shot noise in *F*_*i*_. VMAP measurements were insensitive to illumination intensity for intensities > 15 mW/mm^2^, providing broad robustness to inhomogeneous illumination or intensity drift of the light source (Fig. S2).

### Validation of VMAP

We validated the method via whole-cell voltage clamp recordings in HEK293 cells expressing Voltron2-JF608, imaged in a customized spinning disk confocal microscope (*24*) equipped with a 607 nm laser (intensity: 100 mW/mm^2^) and an acousto-optic tunable filter for gating the illumination (Fig. 1B, S3A; **Methods**). The fluorescence ratio *R*_v_ was calculated pixel-wise and visualized as spatial maps (Fig 1C). Across *n* = 10 cells, *F*_i_ showed ∼8-fold cell-to-cell variation in basal brightness, but no detectable dependence on V_m_, confirming that the short pulses selectively probed the voltage-insensitive state (Fig. 1D). As expected for Voltron2, *F*_*v*_ decreased monotonically with increasing V_m_, but raw *F*_*v*_ values also varied ∼8-fold between cells (Fig. 1E). The fluorescence ratio *R*_*v*_ = *F*_*v*_/*F*_*i*_ collapsed all cells onto a shared voltage-dependent curve (Fig. 1F). We then performed patch clamp calibration of VMAP in primary neurons (*n* = 7 neurons) and human induced pluripotent stem cells (*n* = 8 cells). The three cell types yielded overlapping curves for *R*_*v*_ vs. V_m_, indicating that the VMAP calibration generalizes across cell types (Fig. 1G; Fig. S4). Over the voltage range from –90 mV to –10 mV, *R*_v_ exhibited an approximately linear dependence on V_m_, described by *R*_v_ = –0.0027 V_m_ + 1.172 (where V_m_ is measured in mV and the data are pooled across cell types). The variability in *R*_v_ across all cells of the three cell types was σ(*R*_v_) = 0.018, corresponding to an absolute accuracy of 6.7 mV (Fig. 1G).

We then co-expressed Voltron2-JF608 and the inward-rectifier potassium channel K_ir_2.1 in HEK cells and imaged the cells on the spinning disk microscope. We varied the extracellular K^+^ concentration to set the resting potential (Fig. 1H) and used VMAP to probe V_m_ of ∼200 cells at each K^+^ concentration. Whereas the voltage-dependent changes in raw fluorescence were obscured by cell-to-cell variations in reporter expression (Fig. 1I, K), the VMAP measurements showed the expected trend with extracellular [K^+^] (Fig. 1J, L). As [K^+^] increased from 2.5 to 90 mM, *R*_v_ changed from 1.480 ± 0.029 (mean ± s.d.) to 1.196 ± 0.021. Based on the calibration by patch clamp, this corresponds to -114 ± 11 mV to -9 ± 8 mV, matching well with the predicted change in K^+^ Nernst potential from -106 mV to -11 mV (assuming intracellular [K^+^] = 140 mM).

Finally, we characterized the molecular mechanism by which Voltron2 switched between voltage-sensitive and insensitive states. Voltron2 is engineered from *Acetabularia* rhodopsin, a light-driven proton pump. Mutations introduced during protein engineering eliminated the steady-state photocurrent (*22*). We reasoned that nonetheless there should be a transient photocurrent upon red light onset, associated with the establishment of a voltage-dependent charge redistribution. In HEK cells expressing Voltron2-JF608, patch clamp measurements showed a transient inward photocurrent when the light turned on, and a transient outward photocurrent when the light turned off (Fig. S5). The sign of these photocurrents is consistent with light-triggered release of a proton into the intracellular space, and then re-uptake during relaxation to the dark-adapted state, in accord with the proposed mechanism of voltage sensitivity in microbial rhodopsin proton pumps (*20*).

### VMAP probes resting potential in cultured neurons

Shifts in neuronal resting potential play an important role in modulating neuronal excitability and spontaneous activity (*25, 26*). Although voltage imaging is widely used to measure neural spike rates and waveforms, it is not typically feasible to image slow shifts in the resting potential. We reasoned that VMAP, combined with blue light activation of a channelrhodopsin, could provide a useful approach for probing neuronal excitability and resting membrane potential.

We first examined whether blue co-illumination affected VMAP measurements. Voltron2-JF608 fluorescence was measured in HEK cells, in the presence or absence of 488 nm co-illumination at 3 mW/mm^2^, an intensity sufficient to saturate activation of the blue light-activated channelrhodopsin CheRiff (*I*_50_ = 0.25 mW/mm^2^ (*27*)). Steady-state blue illumination introduced a slight voltage sensitivity (∼2.8% per 100 mV) into *F*_*i*_, indicating that the blue light partially populated the voltage-sensitive state. In contrast, when combined with steady-state orange illumination, blue illumination changed *F*_*v*_ by less than 1%, presumably because the transition to the voltage-sensitive state was already saturated (Fig. S6). Direct blue light excitation of JF608 fluorescence was negligible. We therefore adopted a protocol where blue light was only applied during the *F*_*v*_ measurements, and not during the dark intervals or *F*_*i*_ measurements.

We then applied VMAP together with optogenetic stimulation to primary rat hippocampal neurons transduced with a lentiviral “Optopatch” construct co-expressing Voltron2 and CheRiff (Fig. 2A; **Methods**). We employed a home-built ultra-widefield microscope equipped with a low-magnification (2×), high-numerical aperture (NA 0.5) objective (*28, 29*). Both red and blue illumination were provided by LEDs. A digital micromirror device (DMD) gated the excitation light for the VMAP protocol (**Methods**). A blue-light ramp followed by step pulses was used to stimulate the neurons (Fig. S3C). At resting membrane potential, VMAP reported an *R*_v_ value of 1.346 ± 0.014 (mean ± s.d., *n* = 218 neurons), corresponding to -64 mV ± 5 mV. Under blue light stimulation, the neurons depolarized progressively, reaching a maximally depolarized baseline *R*_*v*_ of 1.287 ± 0.018, corresponding to -43 mV ± 7 mV. These values are consistent with the expected resting and optogenetically depolarized membrane potentials of neurons (Fig. 2B).

**Figure 2.**
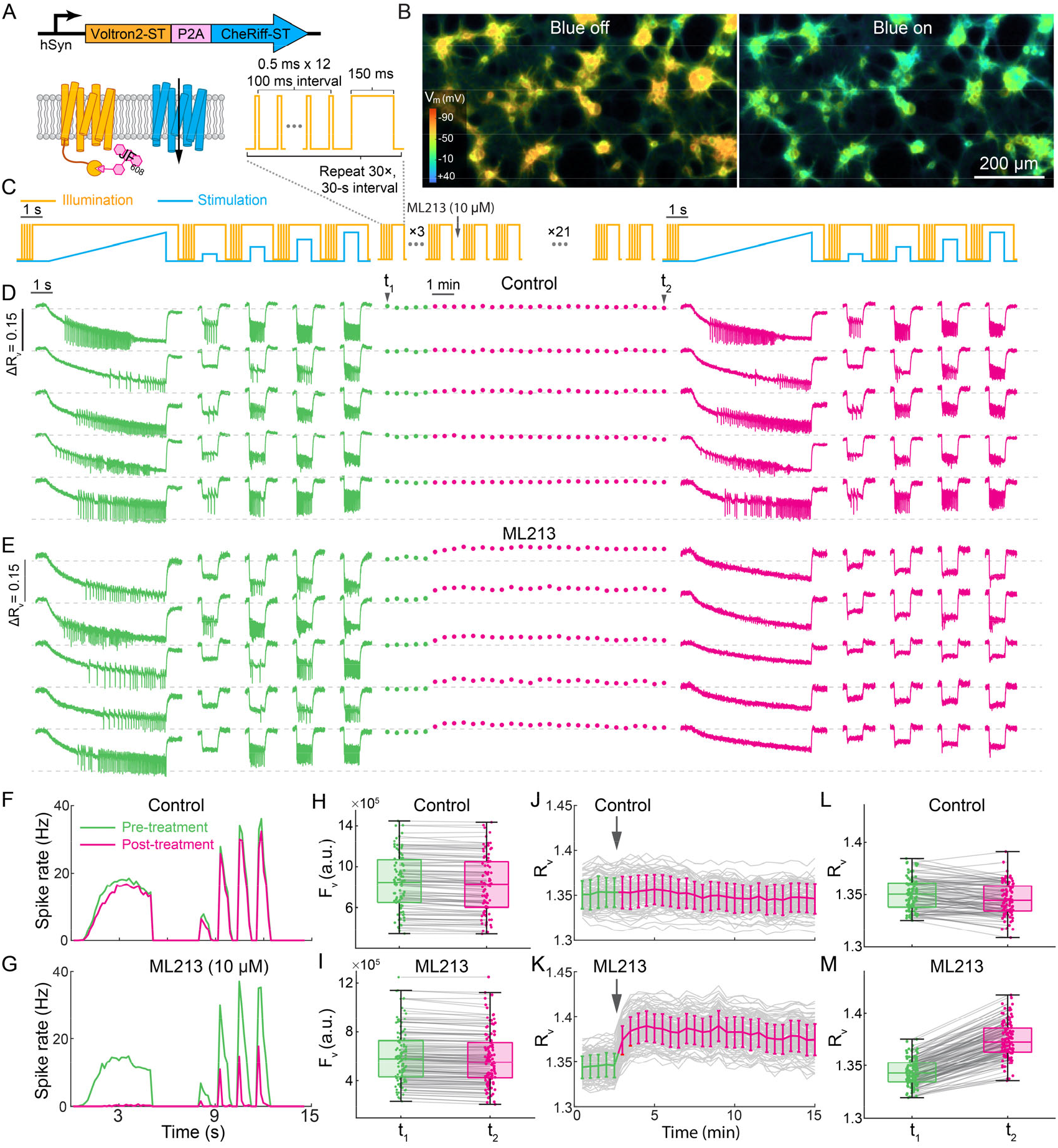
VMAP reveals drug-induced shifts in neuronal resting potential. (A) Optopatch construct for co-expression of Voltron2 and CheRiff, and illumination protocol for shot-noise-robust measurement of *F*_*i*_, followed by *F*_*v*_. (B) VMAP showing change in resting potential of cultured neurons +/- optogenetic stimulation. (C) Experimental protocol combining VMAP measurements with optogenetic stimulation. (D–E) Representative *R*_v_ traces from (D) control and neurons treated with ML213 (10 µM). All the traces show baselines normalized around *R*_*v*_ ∼1.35 (gray dashed lines). (F–G) Optogenetically evoked spike rates before (t_1_) and after (t_2_) neurons were treated with (F) control or (G) ML213. (H–I) Distribution of single-cell fluorescence (*F*_v_) at resting potential before and after the experiment for (H) control and (I) ML213-treated neurons. (J–K) Time course of *R*_v_ at resting potential, across repeated VMAP cycles for (J) control and (K) ML213-treated neurons. (L–M) Comparison of *R*_v_ between initial (t_1_) and final (t_2_) measurements for (L) control and (M) ML213-treated neurons. Boxplots indicate the median and interquartile range (IQR); whiskers extend to the most extreme data points within 1.5×IQR of the lower and upper quartiles. All data acquired on the ultra-widefield epifluorescence microscope.

To probe resting potential dynamics, we then performed a series of 30 VMAP imaging cycles at 30-second intervals (Fig. 2C). After the fifth VMAP cycle we added ML213 (10 µM), an opener of the K_V_7.2 and K_V_7.4 potassium channels, while a control dish was not perturbed. A second Optopatch protocol, identical to the first, then probed drug-induced changes in resting potential and excitability. We recorded a total of 120 neurons treated with ML213 and 98 untreated controls. Representative *R*_v_ traces are shown in Fig. 2D, E. In the control neurons, the optogenetically evoked firing was stable across the experiment (Fig. 2F, Video S1). In the treated neurons, ML213 dramatically suppressed excitability (Fig. 2G, Video S2), consistent with its previously reported action (*30*).

The raw fluorescence *F*_*v*_ decreased slightly between the first (t_1_) and last (t_2_) VMAP measurements in both control (-2.9 ± 3.4%, mean ± s.d., *n* = 98) and drug-treated (-3.8 ± 3.4%, mean ± s.d., *n* = 120) neurons, likely due to photobleaching (Fig. 2H, I). In contrast, the ratiometric VMAP readout clearly distinguished the two conditions: *R*_v_ remained stable in control neurons (Fig. 2J, L) and showed a pronounced drug-induced increase in treated neurons (Fig. 2K, M). The increase in *R*_v_ corresponded to a hyperpolarization of the resting potential from -63 ± 5 mV to a minimum of -80 ± 6 mV (mean shift ΔV_m_ = -17 ± 3 mV), consistent with the ML213 mechanism of action. These results demonstrate that VMAP provides a quantitative, drift-resistant approach for linking changes in neuronal excitability to underlying shifts in baseline membrane potential. More broadly, these results demonstrate mapping of absolute voltage over timescales from ∼1 ms to ∼15 min and spatial scales from ∼2 µm to 3 mm, all in a single experiment. This span of spatial and temporal scales is difficult to achieve by other absolute voltage imaging approaches.

### VMAP reports bioelectrical pattern formation in human stem cell colonies

To test the capability of VMAP to report membrane potential over long times and large areas, we applied VMAP to human induced pluripotent stem cells (hiPSCs) undergoing differentiation. Genetic and pharmacological perturbation experiments have suggested that V_m_ drives the exit from pluripotency (*31*), but the underlying spatiotemporal patterns of V_m_ have not been directly visualized.

We used an hiPSC line that contained a stably integrated Optopatch construct comprising Voltron2 and CheRiff (*28*). Drug-gated degrons (*32*) fused to the GEVI and channelrhodopsin blocked expression until activated by trimethoprim (TMP, 10 µM). TMP addition triggered homogeneous and widespread expression of Voltron2. To ensure that the FRET-opsin GEVI was fully loaded with its two chromophores, we added all-trans retinal (2 µM) and JF608-HaloTag ligand (JF608-HTL, 100 nM) 1-2 hours before each imaging session (Fig. 3A).

**Figure 3.**
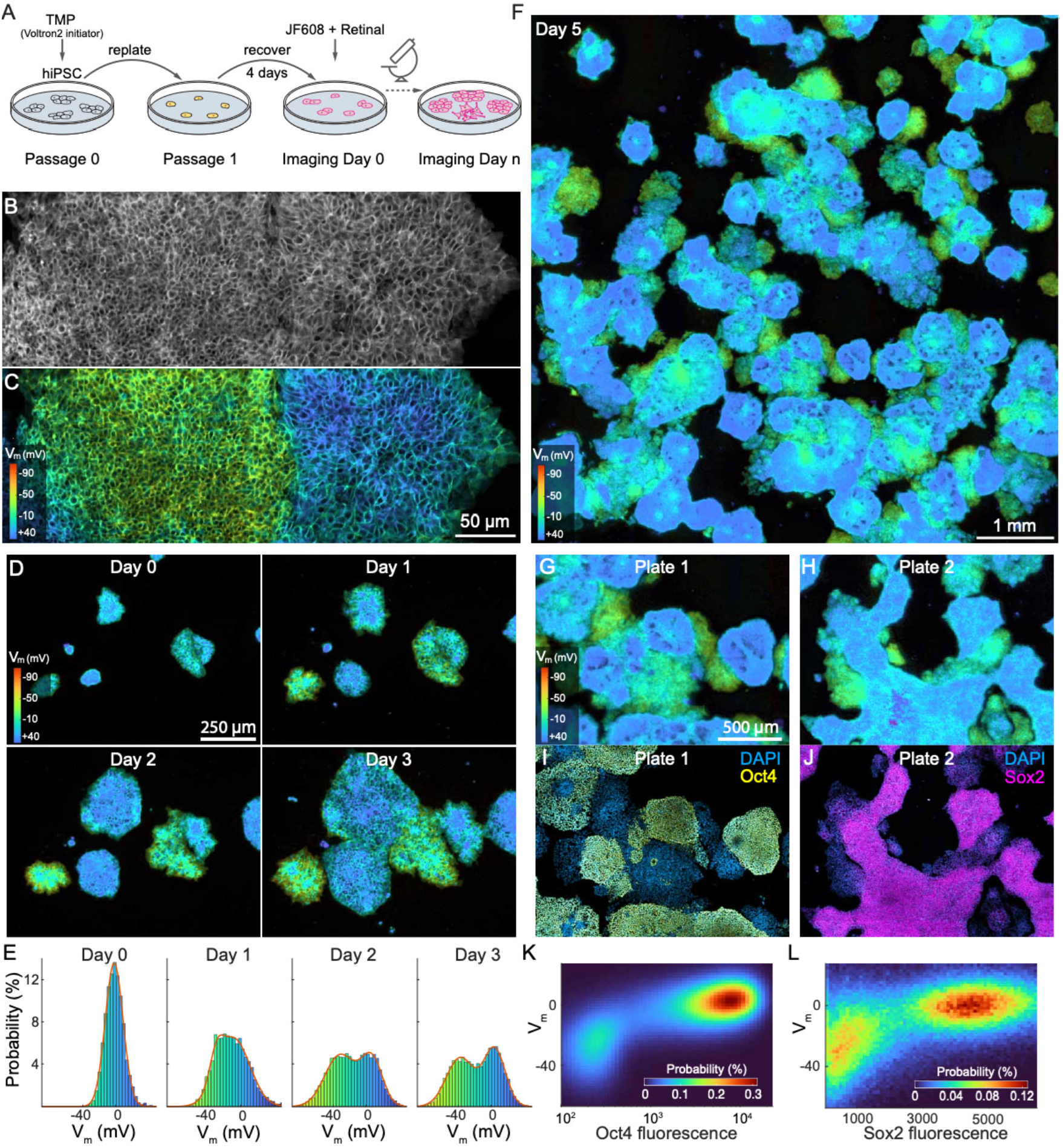
VMAP reports bioelectrical pattern formation in human stem cell colonies. (A) Timeline of hiPSC sample preparation and differentiation. In transgenic Optopatch hiPSC colonies, Voltron2 expression was induced by trimethoprim (TMP). Cells were replated at single-cell density and grown for 4 days, followed by VMAP imaging. (B) High magnification spinning disk image and (C) corresponding *R*_v_ map, revealed spatial domains with distinct membrane potentials (acquired at Day 6). (D) Frames from time-lapse VMAP images of hiPSC colonies over 3 days, showing morphological expansion and emergence of subpopulations with distinct membrane potentials. (E) Distributions of R_v_ on successive days, showing emergence of an electrically polarized sub-population. (F) Wide-area VMAP image demonstrating heterogeneous distributions of *R*_v_ across hiPSC colonies, acquired at Day 6. (G–H) VMAP images of two colonies measured on Day 5. (I–J) Immunofluorescence staining of the same two colonies for pluripotency markers Oct4 and (J) Sox2, showing higher marker expression in depolarized cells. (K–L) Quantification of the relationship between *R*_v_ and marker expression, revealing a strong anticorrelation between membrane potential and expression of (K) Oct4 and (L) Sox2.

We then applied VMAP (waveform in Fig. S3B) to hiPSC colonies, using a spinning disk confocal microscope to achieve high spatial resolution. The raw fluorescence appeared relatively uniform across the colony (Fig. 3B), with cell-to-cell variations in reporter expression dominating any voltage-dependent fluorescence modulation. In contrast, the VMAP images showed clearly resolved multicellular bioelectrical domains with distinct *R*_v_ (Fig. 3C) and sharp boundaries.

We thus sought to map the emergence and dynamics of these domains and to determine their relation to molecular markers of stem cell differentiation. We used the ultra-widefield microscope to perform VMAP imaging on large populations of the hiPSCs. A stage-top incubator maintained physiological conditions for chronic recordings.

VMAP imaging was performed on hiPSC colonies at 30-minute intervals, spanning a 1.8 mm FOV over 72 hours (Fig. 3D, Video S3). Initially, cells formed small clusters with depolarized membrane potentials (*R*_v_ = 1.18 ± 0.03, corresponding to V_m_ = -2 ± 11 mV, mean ± s.d., *n* ∼ 160 cells). Colonies then expanded and segregated into two major populations (Fig. 3D, S7). One population maintained the low *R*_v_ (depolarized) and formed a compact multilayered stack. The other population flattened and exhibited migratory behavior, accompanied by progressively increased *R*_v_ (negative V_m_), reaching 1.26 ± 0.054 (V_m_ = -34 ± 20 mV, mean ± s.d., *n* ∼ 500 cells, Fig. 3E). This shift toward more negative V_m_ during differentiation is consistent with prior reports indicating a relationship between K^+^ channel activation and differentiation (*31, 33–35*).

We next compared these bioelectrical maps with biochemical markers of cell state. We performed VMAP with tiling large FOVs spanning more than 7 mm (∼5×10^4^ cells; Fig. 3F). Following VMAP imaging, hiPSC cultures were fixed and immunostained for the pluripotency markers Oct4 and Sox2. Immunofluorescence images were aligned with the corresponding VMAP images. For both markers, cells with more depolarized membrane potentials exhibited substantially higher marker expression than those with more hyperpolarized V_m_ (Fig. 3G–J, S8). Quantitative analysis revealed two distinct populations with a clear anticorrelation between the fluorescence ratio *R*_v_ and immunostaining intensity for both Oct4 (Fig. 3K) and Sox2 (Fig. 3L). This finding demonstrates that membrane potential, as reported by VMAP, can be used to classify cell state during early differentiation.

### VMAP reveals whole-body bioelectrical patterns in zebrafish embryos

Finally, we applied VMAP (waveform in Fig. S3B) to developing zebrafish embryos to generate whole-embryo membrane potential maps. Voltron2 has previously been used as a voltage indicator in zebrafish heart (*36*), nervous system (*22*), and fins (*37*).

Embryos were injected with Voltron2 mRNA at the single-cell stage, dechorionated, and stained with JF608-HTL for 2 h at ∼17 hours post-fertilization (hpf), followed by a 3-h rinse to remove unreacted dye (Fig. 4A). Imaging at 22-24 hpf was performed using the spinning disk confocal microscope. Using a 25x NA 1.05 objective, we tiled the whole embryo using 40 fields of view, and then acquired a volumetric z-stack with 20 µm inter-plane spacing. Imaging of the whole embryo took ∼25 min over 314 VMAP frames.

**Figure 4.**
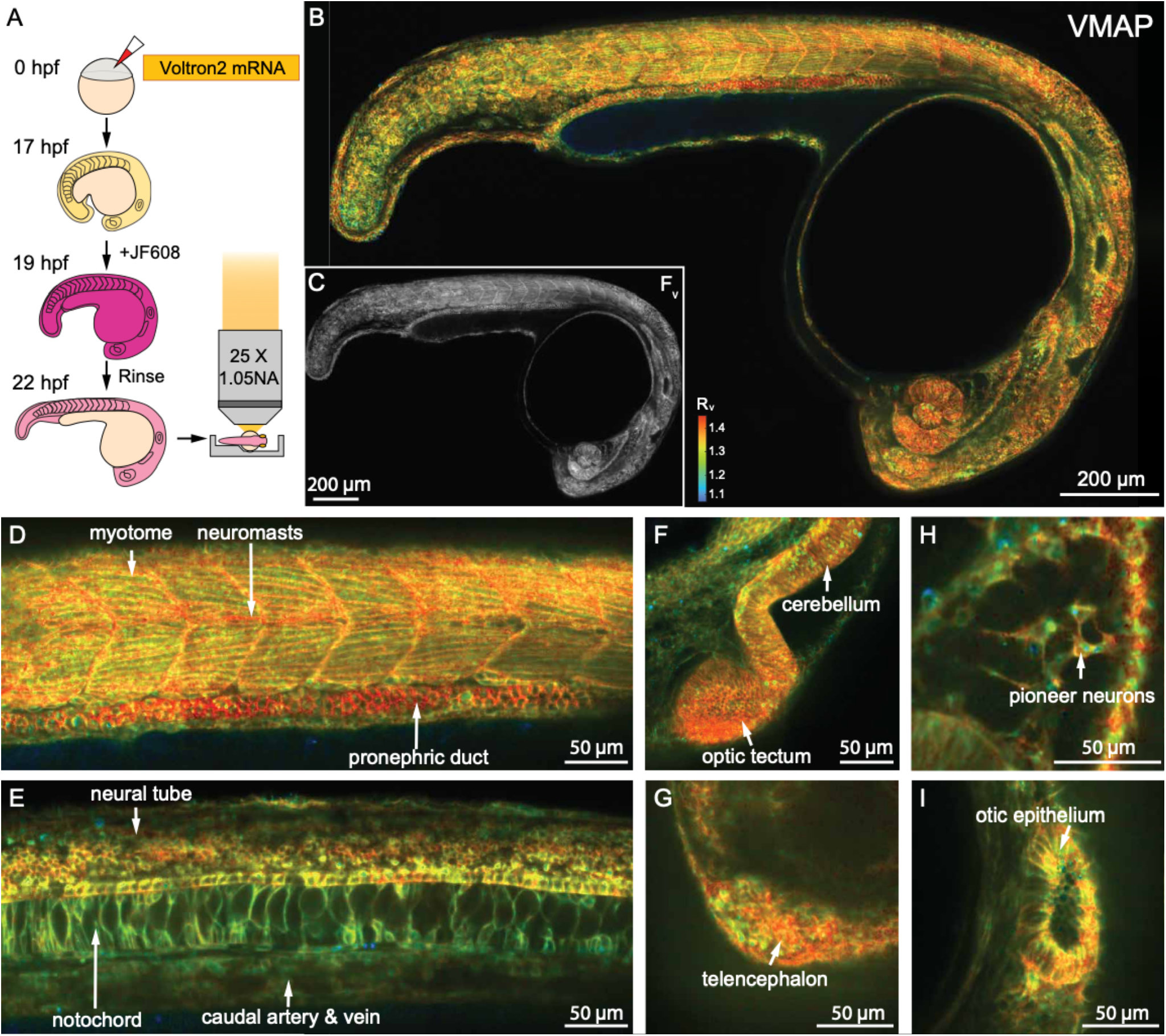
Whole-body voltage maps in zebrafish embryos. (A) Experimental workflow for VMAP imaging in zebrafish embryos. Voltron2 mRNA was injected at the single-cell stage, followed by staining with JF608 and rinsing prior to imaging. Embryos were imaged at 22–24 hpf using a spinning-disk confocal microscope. (B) Maximum intensity projection of a 3D VMAP dataset showing whole-embryo distribution of *R*_v_, revealing spatially heterogeneous membrane potentials. (C) Raw fluorescence, *F*_v_, for the same embryo. (D–E) Enlarged views of the tail region at different depths. (D) Skeletal muscle (myotome) and pronephric duct exhibit elevated *R*_v_, indicative of hyperpolarization. (E) The notochord displays intermediate *R*_v_. (F–G) Brain regions showing strong hyperpolarization, including (F) the optic tectum and cerebellum and (G) the telencephalon. (H) Highly polarized pioneer neurons. (I) Cells in the otic epithelium had comparatively lower *R*_v_ values, indicating a more depolarized state. All data acquired on the spinning disk confocal microscope.

Figure 4B and Video S4 show the result from a 3D VMAP dataset of a whole zebrafish embryo. While the raw fluorescence intensity, *F*_*v*_, appeared mostly uniform (Fig. 4C), the *R*_v_ map revealed striking voltage patterns across the organism. As a negative control, we euthanized the fish (Methods) and then acquired a second 3D VMAP map. In the dead fish we observed a uniform *R*_v_ of ∼1.1, consistent with global membrane depolarization (Fig. S9).

Enlarged views of the live fish’s tail at planes corresponding to skeletal muscle and notochord highlighted distinct bioelectrical states across tissues. Skeletal muscle exhibited higher *R*_v_ values (∼1.37, corresponding to V_m_ ∼ -73 mV), consistent with a more hyperpolarized membrane potential (Fig. 4D), while the notochord showed intermediate values (*R*_v_ ∼1.28, V_m_ ∼ -40 mV, Fig. 4E). The pronephric duct also displayed pronounced hyperpolarization (*R*_v_ ∼1.42, V_m_ ∼ -92 mV, Fig. 4D). In the head, the telencephalon and optic tectum exhibited some of the highest R_v_ values, indicating strongly hyperpolarized potentials (*R*_v_ ∼1.33-1.39 , V_m_ ∼ -59 to -80 mV, Fig. 4F, G), and highly polarized pioneer neurons could be clearly resolved (Fig. 4H). In contrast, cells in the otic epithelium exhibited substantially lower *R*_v_ values (*R*_v_ ∼1.28-1.35, V_m_ ∼ -41 to -66 mV, Fig. 4I), consistent with a more depolarized state. Similar voltage patterns were observed across *n* = 11 fish, measured between 22 and 24 hpf (Fig. S10).

Together, these results demonstrate that VMAP enables volumetric, organism-scale mapping of absolute membrane potential with cellular resolution. The observed spatial heterogeneity in V_m_ across distinct anatomical structures highlights the rich bioelectrical landscape of the developing embryo and underscores the potential of VMAP for studying bioelectric patterning *in vivo* during development.

## Discussion

Because VMAP is independent of any specific imaging modality, the method can be applied flexibly across systems including cell cultures, organoids, and *in vivo*, using the most appropriate imaging platform. We demonstrated VMAP on epifluorescence microscopes, a high-magnification laser-illuminated spinning-disk confocal microscope, and an ultra-widefield microscope with LED illumination. We applied VMAP across rodent, human, and fish; and spanning isolated cells, cultured cell assemblies, and intact animals. We also used VMAP to measure absolute voltage across timescales ranging from millisecond-scale neuronal action potentials to multi-day changes in differentiating hiPSC cultures. Together, these demonstrations establish VMAP as a broadly compatible approach for absolute voltage imaging across diverse experimental contexts.

The VMAP principle can be applied beyond Voltron2-JF608. We also tested Voltron2 labeled with JF585, JF608, or JF635 and illuminated at matched wavelengths, and observed similar illumination-triggered fluorescence activation (Fig. S11). Among other *Acetabularia* rhodopsin– based FRET-opsin GEVIs, Solaris (*38*), but not Positron (*39*), exhibited illumination-triggered activation of voltage sensitivity and was compatible with VMAP (Fig. S12). Thus, VMAP compatibility depends on the photocycle properties of each indicator. Protein engineering of FRET-opsin GEVIs may further tune these photocycles to enhance VMAP performance.

Compared to other absolute voltage imaging techniques, VMAP offers advantages in convenience, throughput, and compatibility. FLIM-based approaches require pulsed lasers, specialized timing electronics, and dedicated detection hardware which are both expensive and difficult to integrate with many existing voltage-imaging platforms (*12–15*). The high photon count needed for precise lifetime measurement also limits the imaging throughput. Two-color ratiometric voltage indicators (*16–19*) requires precise registration of the two imaging channels and are susceptible to artifacts from differential folding, bleaching, degradation or trafficking. The need to image two channels simultaneously also increases hardware complexity and limits the options for multimodal imaging. VMAP avoids these problems by deriving both reference (*F*_*i*_) and signal (*F*_*v*_) from the same fluorophore. Its minimal hardware requirements also facilitate integration with other modalities, such as calcium imaging or optogenetic stimulation (*24, 40*). As with all GEVI-based approaches, VMAP may be affected by confounds that alter voltage sensitivity, including changes in pH, temperature, lipid composition, ionic strength, or reporter trafficking (*9, 13, 17, 41*). In addition, the asynchronous acquisition of *F*_*i*_ and *F*_*v*_ makes VMAP susceptible to fast motion artifacts and creates a trade-off between sufficiently sampling *F*_i_ for accurate normalization vs. preserving time for measuring *F*_*v*_.

VMAP lowers the barrier to investigating a wide range of bioelectrical phenomena. The ability to map absolute voltage in cultured HEK cells (Fig. 1) and neurons (Fig. 2) opens the door to genetic or pharmacological screens – e.g. of ion channel modulators – where resting membrane potential is the readout. The ability to map absolute voltage during differentiation (Fig. 3) and embryonic development (Fig. 4) opens the door to studies on the interplay of bioelectrical, biochemical, and mechanical signaling in models of tissue growth and remodeling, in health and disease. For example, mapping the emergence of membrane potential patterns during embryonic development remains a major challenge for existing techniques, due to the broad spatial and temporal scales involved (*9*). Bioelectrical signals are also proposed to play important roles in wound healing (*37*) and tumor growth (*42*), both of which could be probed by VMAP.

## Acknowledgments

We thank Andrew Preecha, Camila Bodden, and Megan Hershfield for technical assistance. We thank Sharad Ramanathan for guidance on hiPSC culture and immunostaining. We thank Richard She in the Jonathan Weissman Lab for plasmids involved in producing the Optopatch hiPSC stable lines. We thank the Harvard Center for Biological Imaging (RRID: SCR_018673) for infrastructure and support.

## Author contributions

DG developed the VMAP technique, designed and built the ultra-widefield optical system, prepared the HEK cells and zebrafish samples, acquired data, analyzed data, and wrote the manuscript. JZ developed the stable Optopatch hiPSC lines and prepared the neuron and hiPSC samples. MRH measured photo-activation in different opsins. BZJ advised on development of VMAP in zebrafish. XW developed the spinning disk imaging system. DG, JZ, MRH and BZJ developed the genetic constructs. AEC designed the project, oversaw execution, and wrote the manuscript.

## Data, code, and materials availability

VMAP datasets are available at doi.org/10.7910/DVN/YRNOFH and are publicly available as of the date of publication.

## Supplementary Materials

Materials and Methods

Figs. S1 – S12

Tables S1

Movies S1 to S4

Reference

