## Supplementary Information for "Absolute voltage mapping using dynamic photocycle control"

**Supplementary Material**

Materials and Methods
Figures S1 – S12
Table S1
Video captions S1 – S4
References

**Materials and Methods**

*Genetic constructs*

All constructs were generated using standard molecular cloning methods. Briefly, plasmid backbones were linearized by double digestion with restriction enzymes (New England Biolabs) and purified using the GeneJET Gel Extraction Kit (Thermo Fisher Scientific). DNA inserts were generated by PCR amplification and assembled into the linearized backbones using the NEBuilder HiFi DNA Assembly Kit (New England Biolabs). All plasmids were verified by whole-plasmid sequencing (Plasmidsaurus).

For HEK cell experiments (Fig. 1), Voltron2 was expressed from a CMV promoter as CMV::LR-Voltron2 (Addgene plasmid #258172), where LR denotes the membrane trafficking/localization sequence derived from Lucy-Rho (*1*). The $K_{ir}$2.1 was expressed as CMV::$K_{ir}$2.1-mGFP (Addgene plasmid #258170).

For cultured neuron experiments (Fig. 2), Voltron2 was co-expressed with the blue-shifted channelrhodopsin CheRiff using a self-cleaving P2A peptide. The construct consisted of hSyn::Voltron2-ST-P2A-CheRiff-EGFP-ST (Addgene plasmid #258171), where ST denotes the soma-targeted trafficking motif from Kv2.1 that restricts the expression near the soma (*2*).

For hiPSC experiments (Fig. 3), an Optopatch-degron construct, comprising Voltron2-ecDHFR(C12)-ST-P2A-Cheriff-mEOS-ecDHFR(c12)-ST-P2A-mNeon, was integrated into an exon of the β-actin gene via SLEEK method (*3*). The ecDHFR(C12) degron was used to mitigate silencing associated with chronic expression. The template SLEEK donor construct targeting the β-actin locus and the pX458 plasmid containing SpCas9 and an sgRNA targeting the β-actin locus were gifts from Richard She in Jonathan Weissman lab.

For zebrafish embryo experiments (Fig. 4), a construct consisted of Voltron2-KGC-ER2 (Addgene plasmid #258173), was used for *in vitro* mRNA transcription of Voltron2 mRNA. The KGC motif was included to improve membrane localization (*4*), and the ER2 trafficking sequence was included to reduce endoplasmic reticulum retention and improve surface expression (*5*). A construct consisted of alpha-bungarotoxin was used for in vitro mRNA transcription of alpha-bungarotoxin mRNA. The alpha-bungarotoxin plasmid was a gift from Sean Megason (Addgene plasmid #69542).

*HEK cell culture*
HEK293T cells were maintained in tissue culture-treated dishes (Corning) at 37 °C and 5% $CO_2$ in Dulbecco's Modified Eagle Medium supplemented with 10% fetal bovine serum, 1% GlutaMAX-I, penicillin (100 U/mL), and streptomycin (100 μg/mL).

For patch-clamp experiments (Fig. 1B–G), cells in 35-mm dishes were transiently transfected with the imaging construct using Lipofectamine 3000 reagent (Thermo Fisher Scientific). For each transfection, 0.5 μg of DNA was delivered according to the manufacturer's protocol. Cells were replated 36 h after transfection onto poly-D-lysine-coated glass-bottom dishes (Cellvis, Cat. # D35-14-1.5-N) to promote cell adhesion. For experiments examining $K_{ir}2.1$ under different extracellular $K^+$ concentrations (Fig. 1H–L), cells were virally transduced with lentivirus.

*Neuron culture*
Primary E18 rat hippocampal neurons (fresh, never frozen; BrainBits, #SDEHP) were dissociated according to the vendor's protocol and plated on poly-D-lysine- and laminin-coated glass-bottom dishes. Neurons were plated at 21k cells/cm² and co-cultured with primary rat glia plated at 27k cells/cm² to promote neuronal health and maturation. For lentiviral expression, neurons were transduced after 5–10 days in culture with 200 μL of low-titer lentivirus-containing medium. For Optopatch co-expression experiments, neurons were transduced after 5 days in culture with 25 μL of lentivirus containing the Optopatch construct and imaged 3–4 days later.

*Lentiviral transduction*
All the lentivirus preparations were made in house. HEK293T cells were co-transfected with the second-generation packaging plasmid psPAX2 (Addgene #12260), envelope plasmid VSV-G (Addgene #12259) and transfer plasmids at a ratio of 9:4:14. For small batches, 5.6 μg total plasmids for a small culture (300k cells in 35 mm dish) gave sufficient yield of lentivirus. Lentivirus was not further concentrated. For lentiviral transduction, 100 μL of lentivirus was added to a single 35 mm dish. After 48-60 hours, cells were either replated onto glass for imaging or split and replated on 35 mm plastic dishes for continued growth. Virally transduced cultures could be used for up to three passages after transduction. For all experiments, imaging was performed 12-24 hours after replating on glass.

*Establishment and maintenance of Beta-Actin Optopatch iPSC line*
Details of the construction and characterization of the Optopatch iPSC line are given in (*6*) . Briefly, the parental human iPSC line (WTC11), containing a single-copy integration of CAG-dCas9-BFP-KRAB at the CLYBL locus and Tet-On NGN2 at the AAVS1 locus (introduced via TALEN-mediated integration), was established as previously described (*7*). To generate a monoclonal stable iPSC line expressing Optopatch (Voltron2 and CheRiff), we employed SLEEK

(*3*) to integrate the Optopatch construct and an mNeonGreen selection marker into the exon of the housekeeping gene β-actin (ACTB). The Optopatch-Degron SLEEK construct and pX458 targeting the β-actin locus were co-delivered into the parental iPSC line using Lipofectamine™ Stem (Invitrogen). Approximately one week post-transfection, a subpopulation of cells exhibited mNeonGreen fluorescence. Cells were dissociated and sorted by FACS based on green fluorescence, then plated as single cell per well into two 24-well plates in eTeSR™ medium (STEMCELL Technologies). Medium was not changed until colonies derived from single cells became visible. Large colonies were subsequently dissociated and expanded, and multiple monoclonal lines were established from individual wells.

Culture plates were coated overnight with Geltrex (Gibco, A1413301) diluted 1:100 in DMEM/F12 (Gibco, 11320033). Human iPSCs were maintained on Geltrex-coated plates in mTeSR1 medium (STEMCELL Technologies, 85850) at 37°C with 5% $CO_2$. Medium was replaced daily. When cultures reached 80–90% confluency, cells were dissociated with Accutase (STEMCELL Technologies, 7920). Accutase was quenched by dilution 1:5 in mTeSR1, and cells were collected into conical tubes and centrifuged at 300 × g for 5 min. The supernatant was aspirated, and cells were resuspended in mTeSR1 supplemented with 10 µM Y-27632 dihydrochloride ROCK inhibitor (STEMCELL Technologies, 72308). Cells were then replated onto Geltrex-coated plates at a 1:20 split ratio. iPSCs were maintained in mTeSR1 containing 10 µM Y-27632 for the first day after passaging, after which the medium was replaced with standard mTeSR1. For voltage imaging of the Optopatch iPSC stable line, iPSCs were cultured in mTeSR1 supplemented with 10 µM trimethoprim (TMP; Cayman Chemical, 16473) for one passage to induce Optopatch expression. After 3–4 days of continuous TMP induction, confluent iPSCs were passaged onto imaging dishes or plates as required for downstream experiments.

*Electrophysiology*
All whole-cell recordings were performed in voltage-clamp mode. Immediately before imaging, culture medium was removed, and cells were rinsed and then covered with dye-free extracellular (XC) buffer. The XC buffer contained 125 mM NaCl, 2.5 mM KCl, 2 mM $CaCl_2$, 1 mM $MgCl_2$, 15 mM HEPES, and 25 mM glucose. The buffer was adjusted to pH 7.3 with NaOH and to 305–310 mOsm with sucrose, as measured using a vapor-pressure osmometer (Wescor). Filamented patch pipettes were pulled using an automated puller (Sutter P-1000) to a tip resistance of approximately 5 MΩ and filled with intracellular (IC) buffer containing 8 mM NaCl, 130 mM $CsMeSO_3$, 0.5 mM EGTA, 4 mM Mg-ATP, 0.3 mM $Na_3$-GTP, 3 mM QX-314, and 10 mM HEPES, with pH adjusted to 7.2 using CsOH. Whole-cell voltage-clamp recordings were acquired as described previously (*8*). Signals were amplified using a Multiclamp 700B amplifier (Molecular Devices) and digitized at 100 kHz using a National Instruments DAQ system (NI-PCIe-6323).

*Sample preparation for voltage imaging in cultured cells*
The selected JFDye-HaloTag ligand was added directly to the culture medium at a final concentration of 100 nM and incubated for at least 10-30 min. When imaging hiPSCs, all-trans retinal (2 mM) was added at a final concentration of 2 mM and incubated for 1-2 h. Cells were then rinsed three times and returned to dye-free growth medium. Immediately before imaging, cells were rinsed three additional times and transferred into XC buffer. When imaging hiPSCs over days, the cells were transferred to mTeSR1 medium without phenol red (STEMCELL Technologies, #05877).

*Immunostaining*
Cells were rinsed once with PBS, fixed with 4% paraformaldehyde for at least 15 min, rinsed twice with PBS, and quenched with 100 mM $NH_4Cl$ for 20 min. Cells were then washed three times for 5 min each with PBS, permeabilized with 0.5% Triton X-100 in PBS for 10 min, rinsed three times for 1 min each with 0.1% Triton X-100 in PBS, and blocked in 2% BSA for 10 min. Cells were incubated overnight at 4 °C on a shaker with primary antibodies against OCT4 (1:400; Cell Signaling Technology, #2840) or SOX2 (1:400; Invitrogen, #48-1400). Samples were then washed three times for 5 min each with 0.1% Triton X-100 in PBS and incubated for at least 1 h with secondary antibody (Fisher, Donkey anti-Rabbit, AF555, #A-31572) and Hoechst. Cells were washed three times for 5 min each with 0.1% Triton X-100 in PBS, followed by two 5-min washes with PBS. All immunostaining images were acquired on a Zeiss LSM 980 confocal microscope using 405-nm and 561-nm excitation and a 10× C-Apochromat objective (10×/0.45).

*Zebrafish maintenance*
All vertebrate experiments were approved by the Institutional Animal Care and Use Committee of Harvard University. Zebrafish (*Danio rerio*) of the AB wild-type strain were used for all experiments. Adult fish were maintained at 28.5 °C on a 14 h light/10 h dark cycle. Embryos were collected from crosses between female and male adults aged 3–24 months. Embryos were euthanized by immersion in an ice bath for at least 1 h, followed by freezing to ensure death.

*RNA constructs and microinjection*
Synthetic mRNAs were transcribed from linearized pMTB plasmids (*9*) containing the insert of interest using the mMESSAGE mMACHINE SP6 *in vitro* transcription kit (Thermo Fisher Scientific). mRNAs were injected into embryos at the one-cell stage together with 0.1% phenol red.

*Sample preparation for fish*
Embryos were injected at the one-cell stage with 150 pg of Voltron2 mRNA together with 20 pg of mRNA encoding alpha-bungarotoxin, a peptide acetylcholine receptor blocker used to immobilize tail movements (*10*). Embryos were raised at 28.5 °C until 17 h post-fertilization (hpf). Chorions were removed by incubating embryos in 1 mg/mL Pronase protease (Sigma) in 0.3× Danieau buffer for 6 min at 28.5 °C, followed by gentle mechanical agitation. The 0.3× Danieau buffer contained 17.4 mM NaCl, 0.21 mM KCl, 0.12 mM $MgSO_4$, 0.18 mM $Ca(NO_3)_2$, and 1.5 mM HEPES, pH 7.2. Dechorionated embryos were rinsed twice with 0.3× Danieau buffer and then stained with 500 nM JF608 for 2 h at 28.5 °C. Embryos were subsequently washed twice for 5 min each in 0.3× Danieau buffer and incubated for an additional 2 h at 28.5 °C in 0.3× Danieau buffer to remove unbound JF dye. Tail-array mounts were cast in 2% agarose in 0.3× Danieau buffer using a custom mold designed to mount embryos flat in 60-mm Petri dishes (*11*). Embryos were mounted in 0.3x Danieau and affixed by placing a 0.17 mm glass coverslip over the mount array.

*Customized spinning disk microscope*
Figures 1, 3B, 3C, 4, S2, S4, S9 and S10 were imaged on a custom-built upright spinning-disk confocal fluorescence microscope (*12*). Illumination was provided by a 607-nm laser (Opto Engine LLC, MLL-FN-607, 600 mW). Laser intensity was modulated using an acousto-optic tunable filter (AOTF; Gooch and Housego, TF525-250-6-3-GH18A). The laser beam was relayed through a 4f

optical system consisting of two 400-mm convex lenses and coupled into a single-mode optical fiber for spatial homogenization before delivery to the microlens array of a commercial spinning-disk unit (Yokogawa, CSU-X1). The microlens array focused excitation light onto the pinhole array of the spinning-disk unit. An excitation tube lens (Thorlabs TTL100-A) relayed the intermediate image plane to a 25× water-immersion objective with 1.05 numerical aperture (Olympus XLPLN25XWMP2), yielding an effective magnification of 14× and re-imaging the spinning-disk pinhole array onto the sample.

Fluorescence was collected through the same objective and tube lens and re-imaged onto the spinning-disk pinhole array for optical sectioning. Emission light was separated from excitation light using a custom dichroic mirror positioned between the pinhole and microlens arrays (Chroma, zt488/607tpc, 13 × 15 × 0.5 mm), passed through an emission filter (IDEX, FF01-709/167-25), and imaged onto a scientific CMOS camera (Hamamatsu ORCA-Fusion) using two 100-mm lenses. The duty ratio of the spinning disk was 4%, corresponding to an effective exposure of 0.04 ms per pixel during a 1-ms illumination pulse.

Hardware control and data acquisition were performed using Luminos software (*13*). Camera acquisition was synchronized by frame-trigger pulses generated by a National Instruments DAQ board (NI-PCIe-6323), referenced to the 100-kHz camera output clock signal (Hsync). The same timing reference was used to synchronize experimental waveforms controlling the AOTF and shutters. To synchronize camera frames with spinning-disk rotation, a standalone microcontroller board (Arduino Uno Rev3) received the camera clock signal and generated a 1-kHz pulse train. This pulse output was connected to the CSU_SYNC input (pin 12 of the PCR68 connector) of the spinning-disk control unit to trigger and stabilize disk rotation.

Data for Figure S1, Figure S5, Figure S6, Figure S9, Figure S11 and S12 were acquired using widefield variants of the similar microscope platform. In these configurations, the spinning-disk confocal unit was omitted while retaining a similar microscope body, objective configuration, camera-based detection, and acquisition workflow. Imaging conditions for each figure are summarized in Table S1.

*Customized ultra-widefield microscope*

Figures 2, 3D–H, S7, and S8 were imaged on a custom-built ultra-widefield microscope (*6*). Illumination was provided by an amber LED (OSRAM, LE A P2MQ, 613 nm) and a blue LED (Luminus, PT-121-B, 460 nm). Amber and blue excitation light were spectrally filtered using a 600/37-nm filter (IDEX, FF01-600/37-32) and a 457/50-nm filter (IDEX, FF01-457/50-32), respectively. Light from each LED was collected by a two-lens assembly (Thorlabs, LA4148-A and ACL3026U-A) and combined using a dichroic mirror (IDEX, FF580-FDi01).

The combined illumination was passed through a homogenizer within a custom digital micromirror device (DMD) assembly (Digital Light Innovations, 3DLP9000) and coupled to the DMD chip for spatiotemporal modulation. Light from the DMD was collected by a 105-mm camera lens (Sigma, ART 105 mm F1.4 DG HSM) and relayed to a 2× objective with 0.5 numerical aperture (Olympus MVPLAPO 2XC). For all imaging performed on this setup, the LEDs were driven at 5 Amps, producing an illumination intensity of approximately 20 mW/mm$^2$ at the sample plane.

Fluorescence was collected through the same objective and separated from excitation light using a custom dichroic mirror (Chroma, zt488/607tpc, 50 × 72 × 3 mm) and an emission filter (IDEX, AF01-653/47). Emitted fluorescence was imaged onto a scientific CMOS camera (Teledyne Kinetix) using a second 105-mm camera lens (Sigma, ART 105 mm F1.4 DG HSM), yielding an effective magnification of 2.3×. For long-term live-cell imaging, a stage-top incubator (Tokai Hit, INUBG2ATFP-WSKM) with temperature and gas control was used to maintain the culture environment.

Hardware control and data acquisition were performed using Luminos software (*13*). Camera acquisition was synchronized by frame-trigger pulses generated from a National Instruments DAQ system (USB-6353) referenced to the DAQ internal 100-kHz clock. The same clock was used to synchronize experimental waveforms controlling DMD pattern switching and LED drive current.

*VMAP data analysis*
All data analysis was performed in MATLAB.

VMAP videos were analyzed by separating fluorescence acquired during short initial probe pulses, $F_i$, from fluorescence acquired during longer voltage-reporting pulses, $F_v$. Illumination timing recorded by the DAQ was first aligned to the camera movie by cross-correlation with the mean fluorescence trace. Short and long pulses were then classified by pulse duration.

A dark frame was calculated for each video by averaging frames preceding the short pulses, typically from 30 ms to 3 ms before pulse onset. This dark frame was subtracted from the full movie to set the fluorescence baseline to zero. For spinning-disk recordings, an additional sector correction (*12*) was applied to remove periodic artifacts from the Nipkow disk. Artifact patterns were estimated from frames acquired during long pulses, when illumination was present throughout the camera exposure, and the resulting phase-dependent correction was applied to all movie frames.

Because fluorescence from a brief $F_i$ pulse could be distributed across multiple camera frames depending on pulse timing relative to the camera exposure, candidate frames within a five-frame window around each short pulse were examined. For widefield imaging, frames were classified as containing $F_i$ signal when their mean intensity exceeded three times the dark-frame noise. For spinning-disk imaging, frames were selected based on strong vertical autocorrelation associated with the spinning-disk illumination pattern, and the brightest 3 qualifying frames were retained. A full $F_i$ image for each short pulse was generated by summing all frames assigned to that pulse.

Frames containing $F_v$ were selected from the plateau phase of each long pulse, beginning 30 ms after pulse-on and ending 3 ms before pulse-off. For single-image VMAP measurements, $F_v$ was averaged across the selected long-pulse frames. For time-resolved VMAP measurements, the selected $F_v$ frames were retained for calculating $R_v$ traces.

When motion was present, movies were corrected using NoRMCorre-based registration (*14*). The mean $F_v$ image was used as the registration reference. For $F_i$, registration was performed on the summed full-$F_i$ images, and the resulting shifts were applied back to the individual frames contributing to each $F_i$ measurement. For $F_v$, motion correction was applied directly to the long-pulse frames.

For recordings with long illumination epochs in which photobleaching was appreciable, $F_v$ trace segments were concatenated and baseline regions without blue-light stimulation were fit with a two-component exponential decay. The estimated bleaching baseline was then segmented, normalized, and used to correct each $F_v$ segment. When multiple short pulses preceded a long pulse, the corresponding $F_i$ measurements were averaged to improve precision.

For snapshot measurements, a single $F_v$ image averaged over the long pulse was divided by the corresponding averaged $F_i$ image. For time-resolved measurements, each $F_v$ frame or trace point was divided by $F_i$.

The VMAP ratio, $R_v$, was calculated as $F_v/F_i$. $R_v$ was calculated either pixel-wise or from user-defined regions of interest (ROIs), depending on the analysis. For VMAP image generation, $R_v$ was calculated independently for each pixel as $F_v/F_i$. This spatial map of $R_v$ was subsequently smoothed by a 3x3 median filter to exclude the possible extreme ratio values. For single-cell or cell-cluster measurements, ROIs were manually drawn around the structure of interest, and a single $R_v$ value was calculated as the ratio of the mean fluorescence values within the ROI, mean($F_v$)/mean($F_i$). When the camera exposure time exceeded the short-pulse duration, the ratio was additionally normalized by the exposure-time to short-pulse-duration ratio.

Pseudocolor VMAP overlays were generated by mapping the $R_v$ ratio image to RGB via the turbo colormap, and then modulating the brightness by the corresponding $F_v$ image. Thus the final RGB overlay represents $R_v$ as hue and $F_v$ as brightness. For time-series data, the same procedure was applied independently to each frame.

*Voltage spike detection*

Candidate spikes in cultured neurons were first identified as local maxima (after the fluorescence trace was inverted) exceeding 8× baseline noise, with minimum temporal separation criteria to remove closely spaced duplicate detections. Peri-event waveforms were extracted, and spike onset was defined as the point of maximal second derivative preceding the peak. Onset-aligned candidate waveforms were averaged to generate a cell-specific spike template.

This template was then used to convolve the high-pass-filtered trace, enhancing events with spike-like kinetics. Final spikes were detected from the convolved trace using a 7× noise threshold and a minimum peak distance of 5 frames. Spike times were refined by locating the local maximum in the original normalized trace within ±2 frames of each detected event. Events too close to the trace boundaries were excluded.

For the data analysis in Figure 2, only neuronal traces with more than 10 spikes in the first round of Optopatch stimulation were included for data analysis.

*VMAP and immunostaining registration*

VMAP images were used as reference. Image registration was performed in MATLAB using a correlation-based approach (imregcorr) to estimate the geometric transformation between the

DAPI image and the VMAP image. The estimated transformation was then applied to align the other immunostained images (Oct4 and Sox2).

*Image collage generation*

Large-field VMAP images were generated by stitching sequentially acquired sub-images tiling the large FOV. The fluorescence intensity image stack and corresponding ratio image stack were first transposed to match the physical scan orientation and cropped to remove edge artifacts. Tiles were then assigned to scan rows according to the acquisition order.

Tile registration was first performed within each scan row using the fluorescence intensity images. Adjacent tiles were aligned by translational registration of their overlapping regions using phase-correlation-based image registration. Specifically, the right-side overlap of each tile was registered to the left-side overlap of the next tile, and the resulting translations were accumulated to determine the relative position of each tile within the row. Each row was then assembled into a provisional row image.

The provisional row images were subsequently aligned to one another by translational registration, again using the fluorescence intensity channel. The resulting row offsets were combined with the within-row tile offsets to generate a global position for every tile in the scan. These registration coordinates, determined from the intensity images, were then applied identically to both the fluorescence intensity stack and the VMAP ratio stack.

Final whole-field images were generated by placing each tile into a common canvas at its registered position. In regions where multiple tiles overlapped, pixel values were averaged using an occupation map that recorded the number of contributing tiles per pixel. This produced a stitched fluorescence intensity image and a stitched VMAP ratio image with averaged overlap regions.

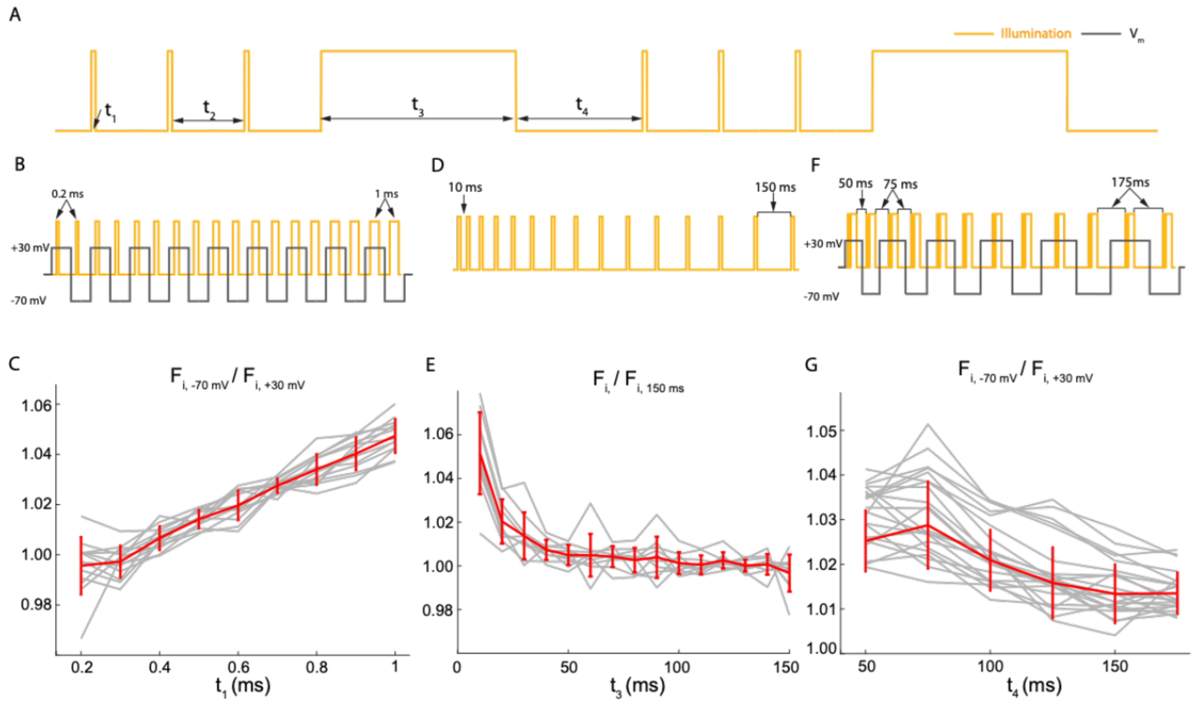


**Figure S1: Optimization of illumination timing for VMAP.** Illumination waveforms were optimized in HEK cells expressing Voltron2-JF608, under voltage-clamp control and 594 nm illumination. (A) VMAP illumination waveform with time intervals marked. $t_1$: short pulse duration. $t_2$: time between consecutive short pulses. $t_3$: long pulse duration. $t_4$: dark time between end of long pulse and subsequent short pulse. The time $t_3$ can be flexibly adjusted (provided that $t_3 >$ 30 ms) to set the measurement cadence and the balance between reference ($F_i$) and voltage ($F_v$) measurements. (B) Waveform to optimize $t_1$: $F_i$ values were successively measured at −70 mV and +30 mV, with short pulse $t_1$ varied from 0.2 to 1 ms. (C) Ratio of $F_i$ at -70 mV to $F_i$ at +30 mV as a function of short pulse duration ($n$ = 13 cells). (D) Waveform to optimize $t_2$, the waiting time between successive short pulses. $F_i$ values were measured with dark intervals from 10 to 150 ms. (E) Ratio of $F_i$ to $F_i$ measured with $t_2$ = 150 ms ($n$ = 10 cells). (F) Waveform to optimize $t_4$, the waiting time in the dark for the GEVI to relax back to a voltage-insensitive state. $F_i$ values were measured with dark intervals from 50 ms to 175 ms. (G) Ratio of $F_i$ at -70 mV to $F_i$ at +30 mV with different dark time ($n$ = 21 cells).

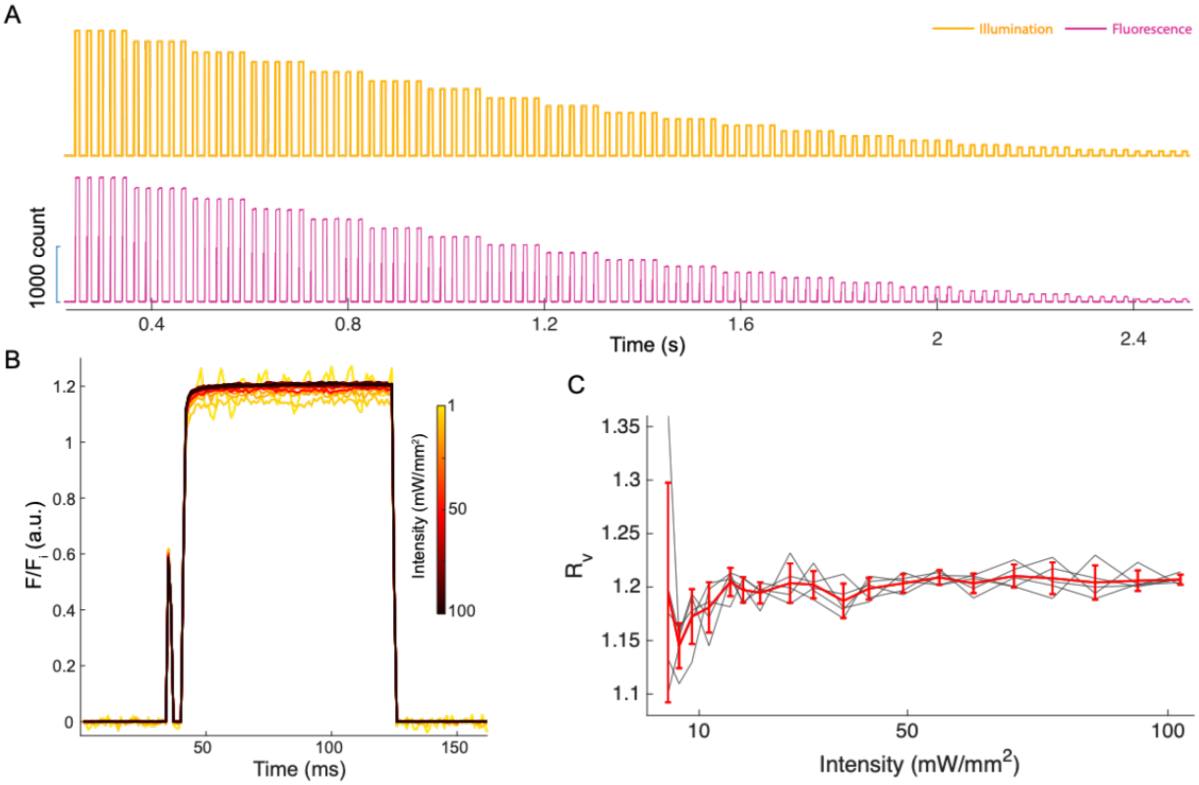


**Figure S2: Illumination intensity required to activate the voltage-sensitive state of Voltron2 for VMAP.** (A) Illumination waveform and fluorescence trace used to test the dependence of VMAP on excitation intensity of 607 nm excitation. (B) Fluorescence responses normalized to $F_i$ at different illumination intensities, showing saturated population of the voltage-sensitive state. (C) $R_v$ vs. illumination intensity. $R_v$ increased with excitation intensity and approached a plateau at intensities > 15 mW/mm², establishing a minimum illumination intensity for VMAP with Voltron2-JF608.

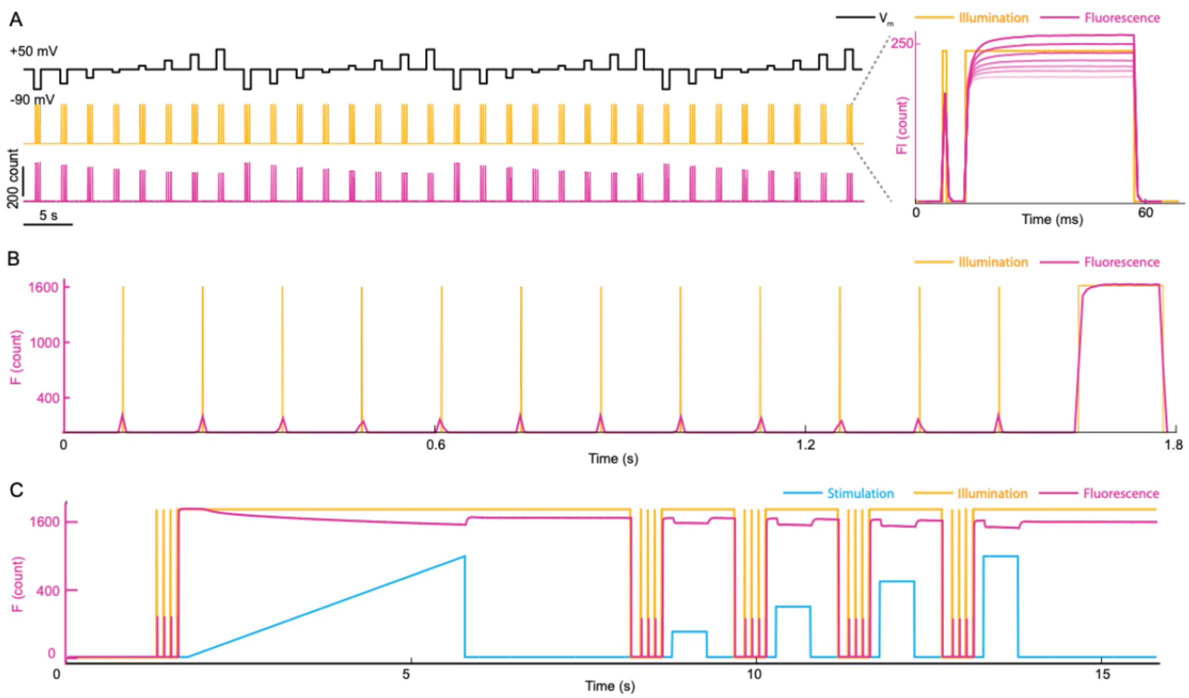


**Figure S3: Acquisition waveforms and fluorescence traces for VMAP measurements.** (A) Complete voltage-clamp VMAP waveform and representative fluorescence response, for the HEK293 patch-clamp calibration experiments in Fig. 1B. (B) Full waveform for acquisition of a shot-noise-robust single VMAP measurement ($R_v$), together with a representative fluorescence trace. The protocol includes 12 short probe pulses, whose fluorescence signals are averaged to obtain a more precise estimate of $F_i$. This protocol was used for the extracellular $K^+$ titration experiments in Fig. 1I, the repeated VMAP sampling in Fig. 2, and all VMAP measurements in Figs. 3 and 4. (C) Complete Optopatch waveform used in the experiments in Fig. 2 (in primary neurons). The fluorescence trace was averaged from the whole field of view.

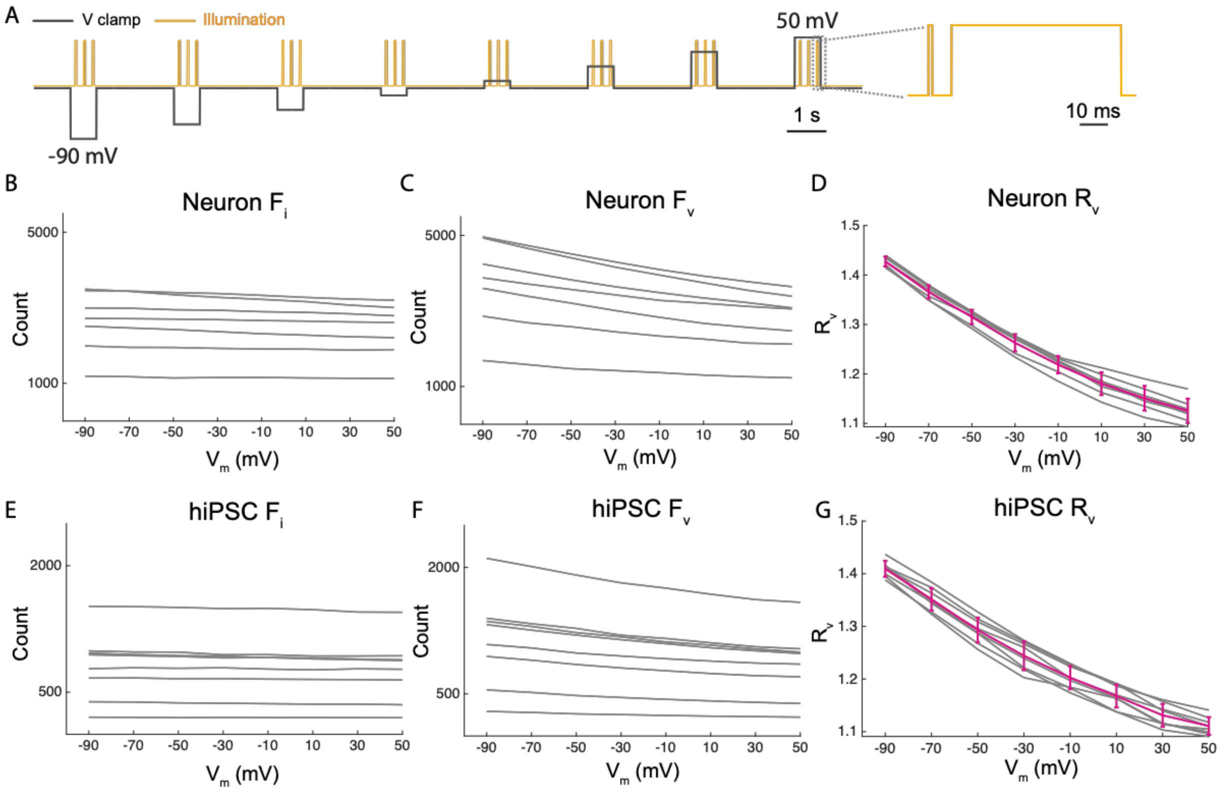


**Figure S4: Patch-clamp calibration of VMAP in cultured neurons and hiPSCs.** (A) Voltage-clamp and illumination waveforms used to calibrate VMAP in cultured cells. (B) Fluorescence measured in the voltage-insensitive state ($F_i$) and (C) voltage-sensitive state ($F_v$) as a function of membrane potential for $n$ = 7 neurons. (D) Calibration curve of $R_v$ vs. holding potential of neurons. $R_v = -0.0026\ V_m + 1.185$. (E) Fluorescence measured in the voltage-insensitive state ($F_i$) and (F) voltage-sensitive state ($F_v$) as a function of membrane potential for $n$ = 8 hiPSCs. (G) Calibration curve of $R_v$ vs. holding potential of hiPSCs. $R_v = -0.0026\ V_m + 1.170$.

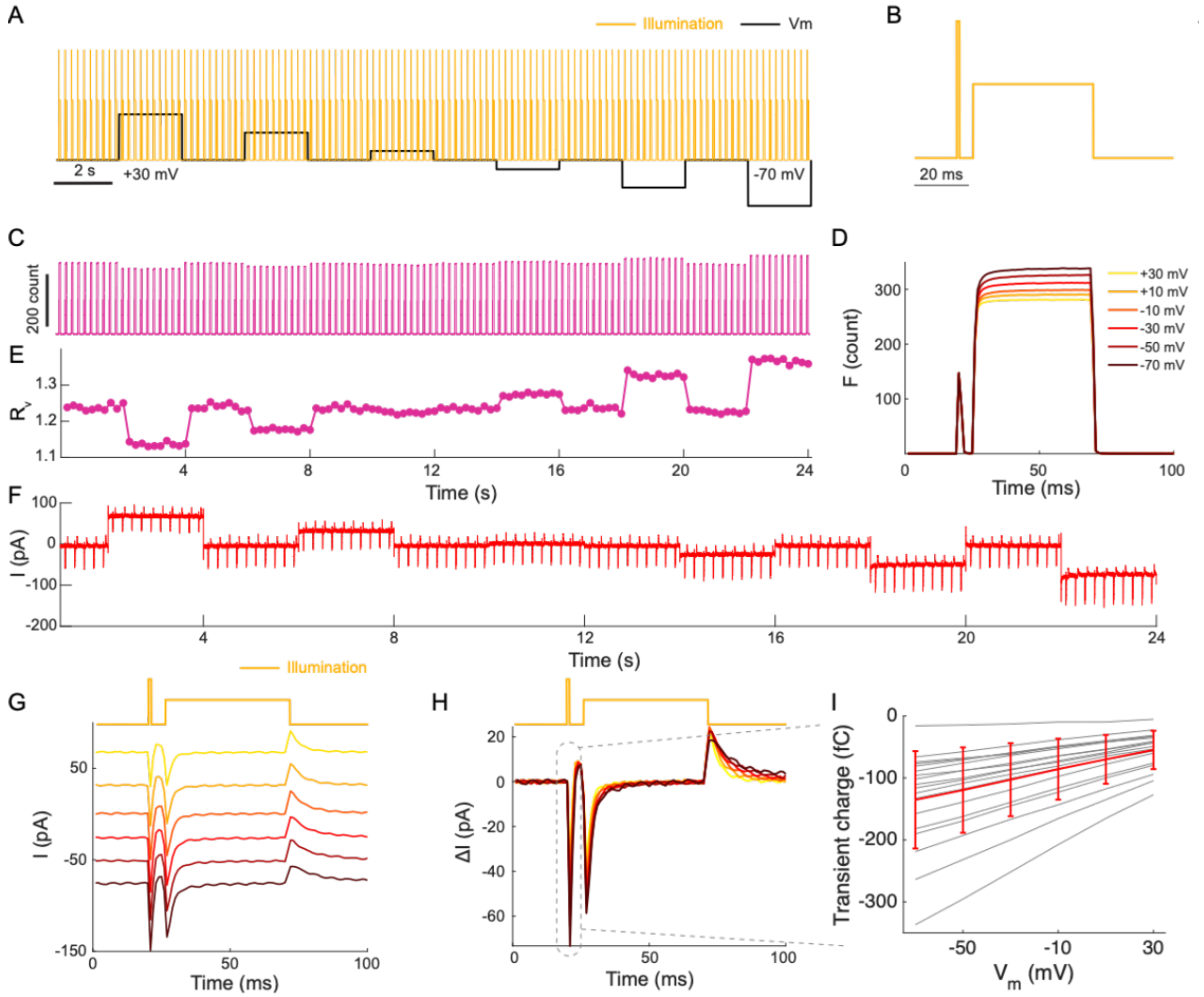


**Figure S5: Transient photocurrents reveal the VMAP mechanism.** (A–B) Voltage-clamp and illumination waveforms used to characterize transient photocurrents. Cells were held at different membrane potentials while illuminated with a modified VMAP sequence in which the short probe pulse was higher intensity than the long pulse. This modification allowed $F_i$ to be measured with sufficient signal from a single short pulse. (C–E) Fluorescence traces recorded under these conditions, and the corresponding VMAP responses. (F) Simultaneously recorded membrane current, revealing a transient inward current upon light-on and a transient outward current upon light-off. (G) Average current traces at different membrane potentials. (H) Baseline-subtracted photocurrents. (I) Peak transient photo-charge as a function of membrane potential, showing an approximately linear dependence on $V_m$. More negative membrane potentials were associated with larger inward photocurrents, consistent with light-triggered establishment of a voltage-dependent equilibrium between protonated and deprotonated Voltron2 Schiff base. We sought to use this photocurrent to estimate the number of Voltron2 molecules per cell. At sufficiently negative $V_m$, we anticipated that each Voltron2 molecule would release $Q_{max} = 1\ e^+$ of charge into the cytoplasm. We did not observe saturation in the photocurrent at negative $V_m$, so we took the absolute value of the photocurrent at $V_m = -70$ mV as a lower-bound. At $V_m = -70$ mV, the integrated photocurrent reached a maximal charge of $|Q| = 135 \pm 78$ fC (mean ± s.d., $n = 18$ cells), corresponding to $Q_{max}/e > 8.1 \pm 4.6 \times 10^5$ Voltron2 molecules per cell (mean ± s.d.).

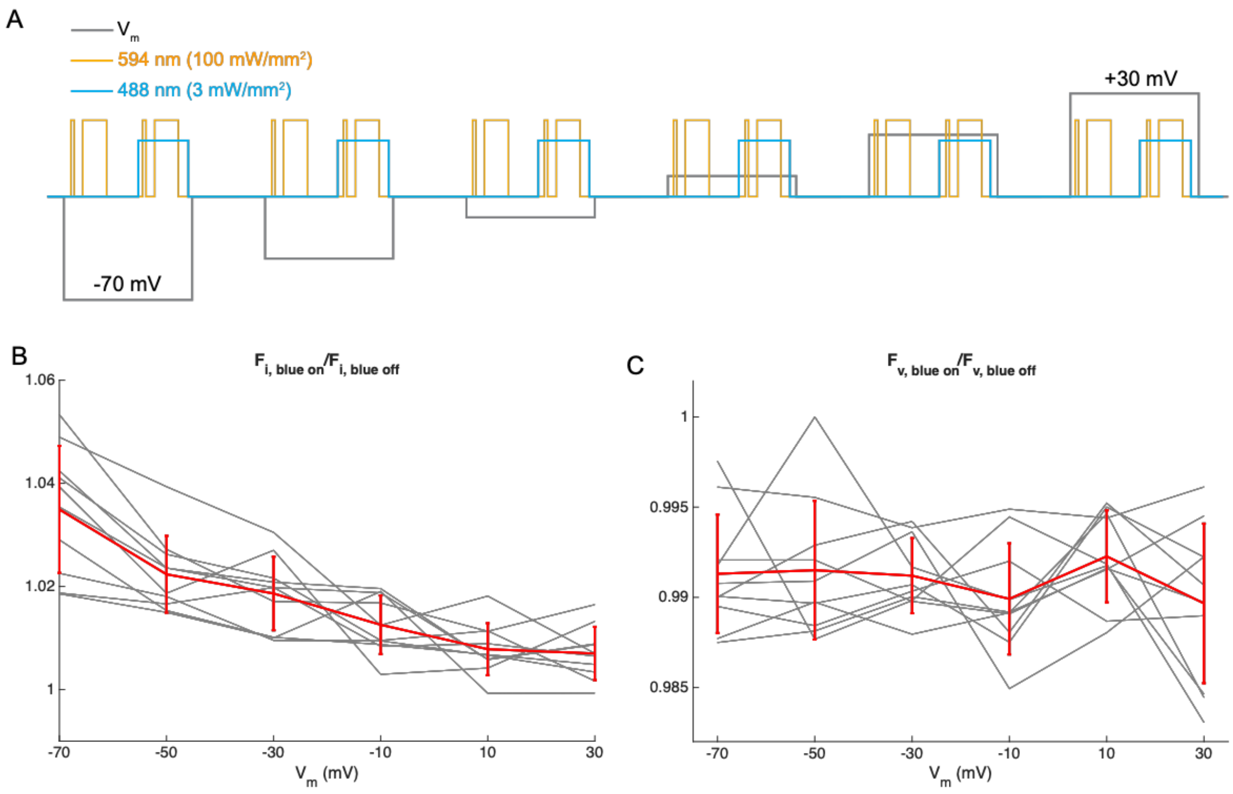


**Figure S6: Effect of blue co-illumination on VMAP fluorescence measurements.** (A) Voltage-clamp and illumination waveforms used to test the effect of 488 nm co-illumination on VMAP, in HEK cells expressing Voltron2-JF608. (B) Ratio of $F_i$ measured with and without blue co-illumination as a function of membrane potential. (C) Ratio of $F_v$ measured with and without blue co-illumination as a function of membrane potential. Across membrane potentials from −70 mV to +30 mV, blue co-illumination increased $F_i$ by ~0.7–3.5%, whereas $F_v$ changed by less than 1%, indicating that VMAP is compatible with blue-light stimulation when blue light is restricted to the measurement period of $F_v$.

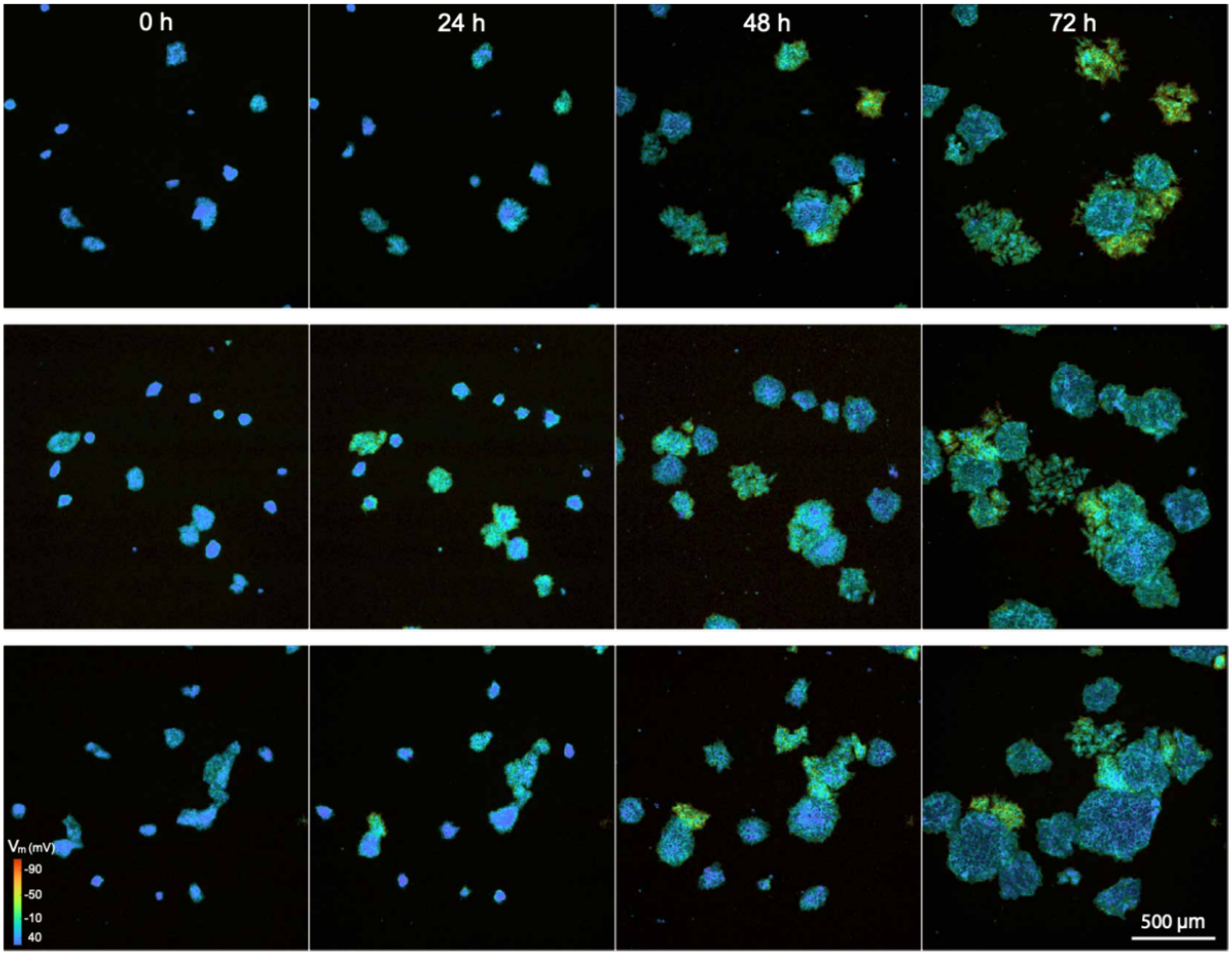


**Figure S7: Additional time-lapse VMAP images during hiPSC differentiation.** VMAP image sequences from three additional hiPSC fields of view acquired over 3 days. As in Fig. 3F, colonies initially appeared as small clusters with relatively depolarized membrane potentials. As differentiation proceeded, the colonies expanded and a subset of cells became progressively more electrically polarized.

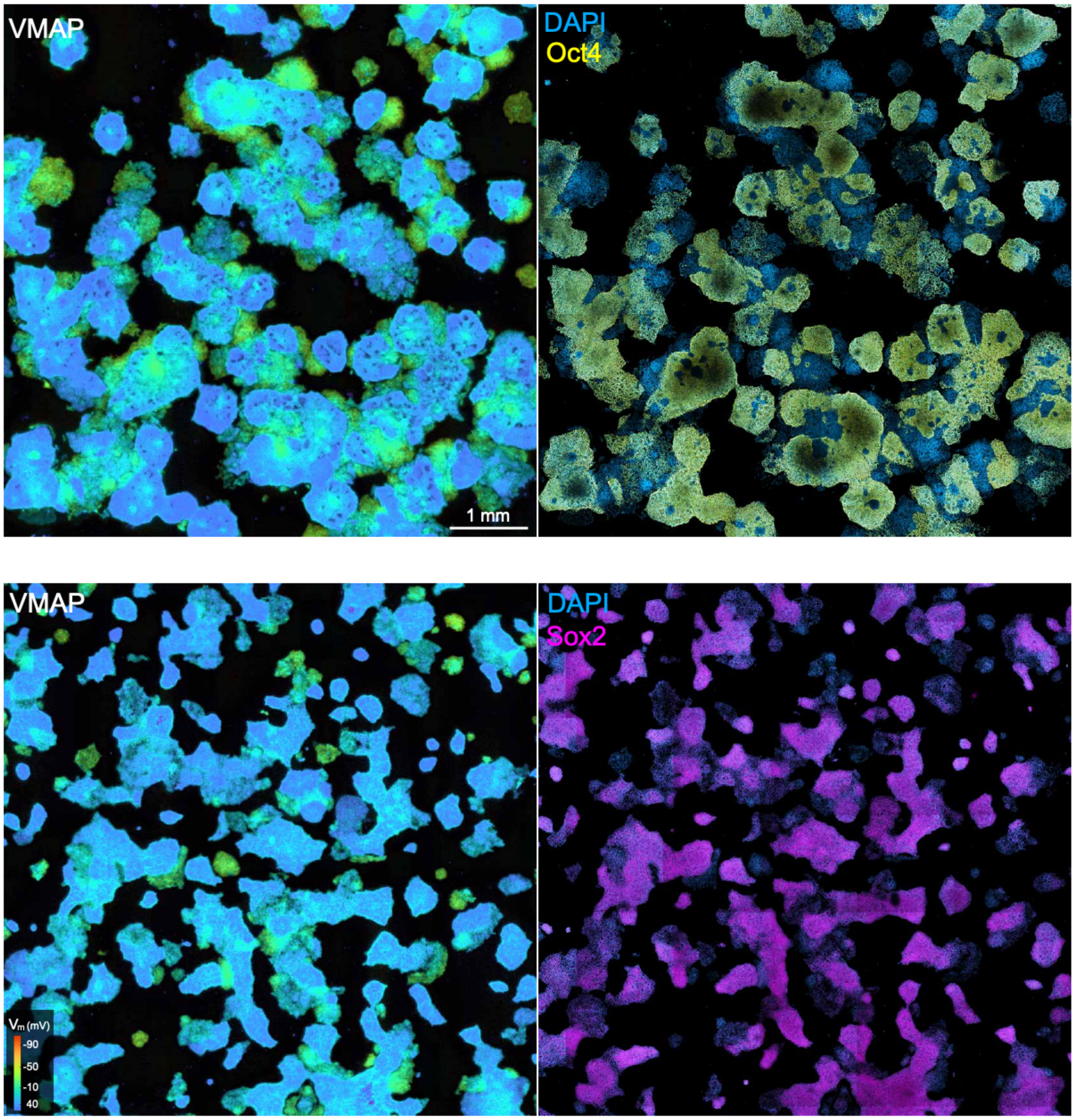


**Figure S8: Wide-area comparisons of VMAP with pluripotency markers in hiPSC cultures.** Full fields of view corresponding to the cropped regions shown in Fig. 3I–L. VMAP images are shown alongside post-fixation immunofluorescence staining of the same fields of view, showing Oct4 or Sox2 and DAPI. In both examples, regions with lower $R_v$ (more depolarized) exhibited stronger Oct4 and Sox2 staining; regions with higher $R_v$ (more polarized) showed little marker expression. These full-field images further illustrate the spatial correspondence between membrane potential and pluripotency in differentiating hiPSC cultures.

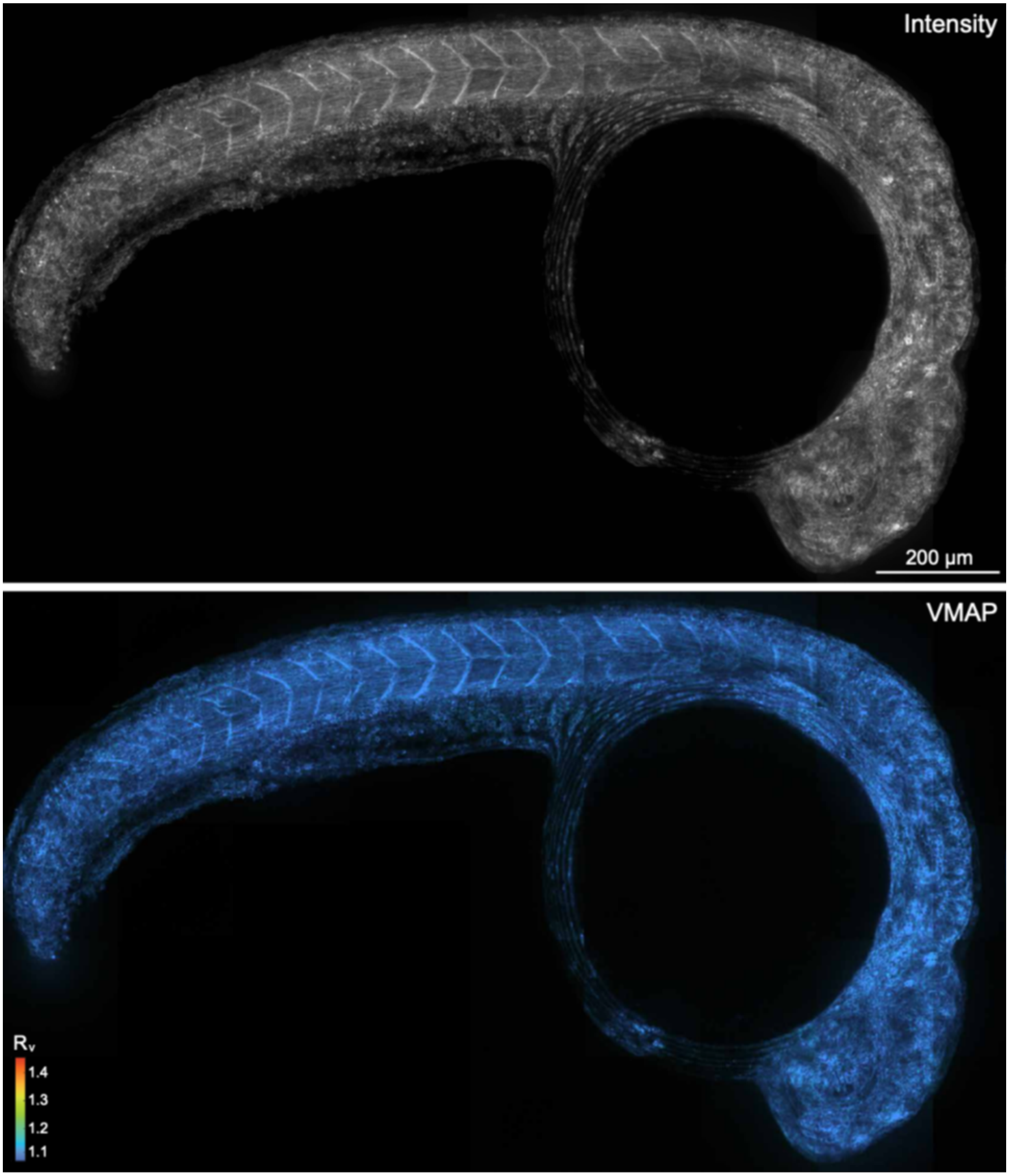


**Figure S9: VMAP map of a euthanized zebrafish embryo.** Maximum-intensity projections of fluorescence intensity (top) and the corresponding VMAP ratio map (bottom) from a zebrafish embryo after euthanasia. In contrast to the spatially heterogeneous $R_v$ patterns observed in live embryos, the dead embryo exhibited a largely uniform $R_v$ of ~1.1 throughout the body, consistent with global membrane depolarization.

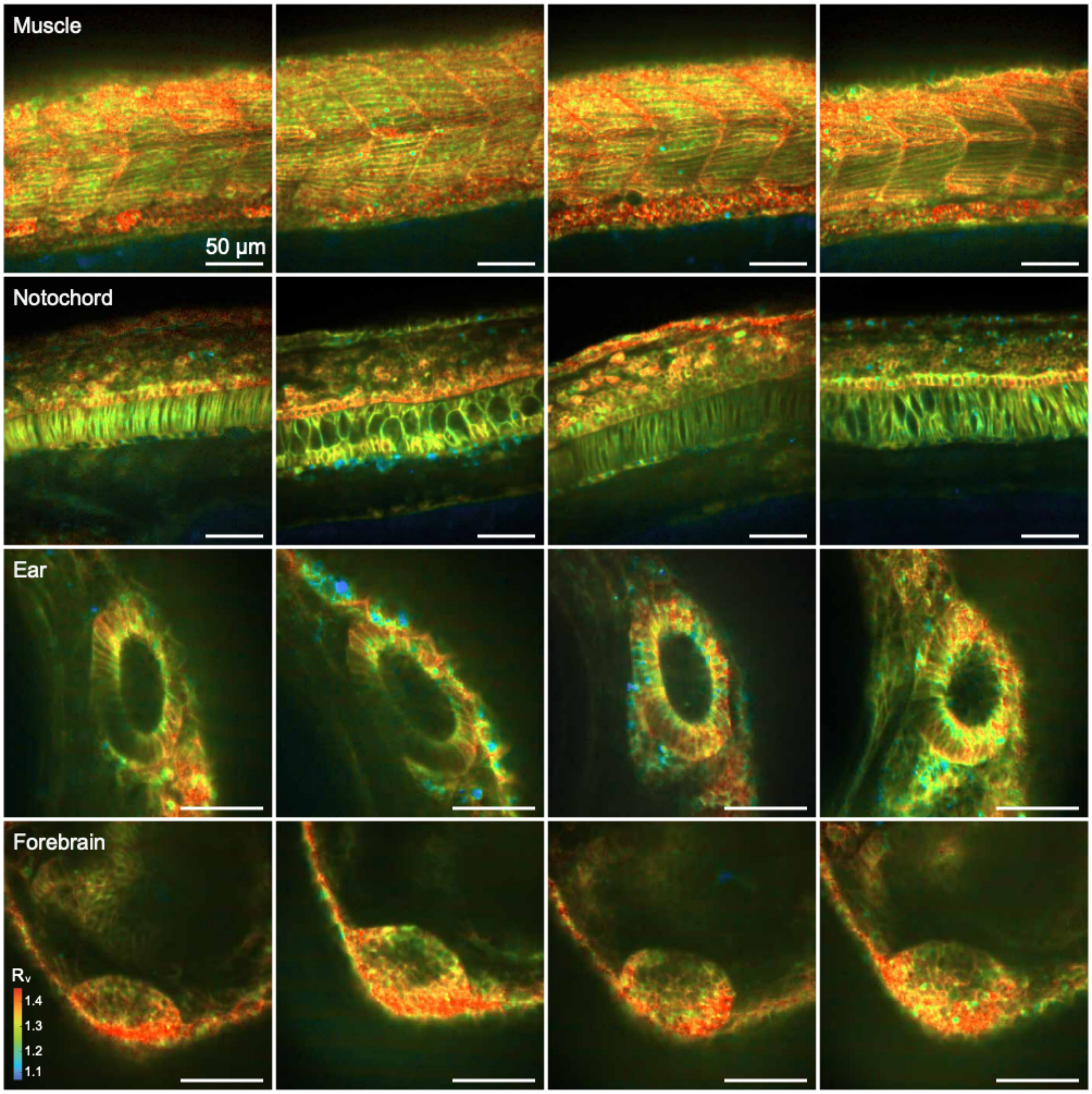


**Figure S10: VMAP measurements in multiple zebrafish embryos.** VMAP measurements from additional zebrafish embryos demonstrate reproducible tissue-specific membrane potential patterns. The data here are obtained from 11 fish (not all regions were imaged in all fish). All scale bars 50 $\mu$m.

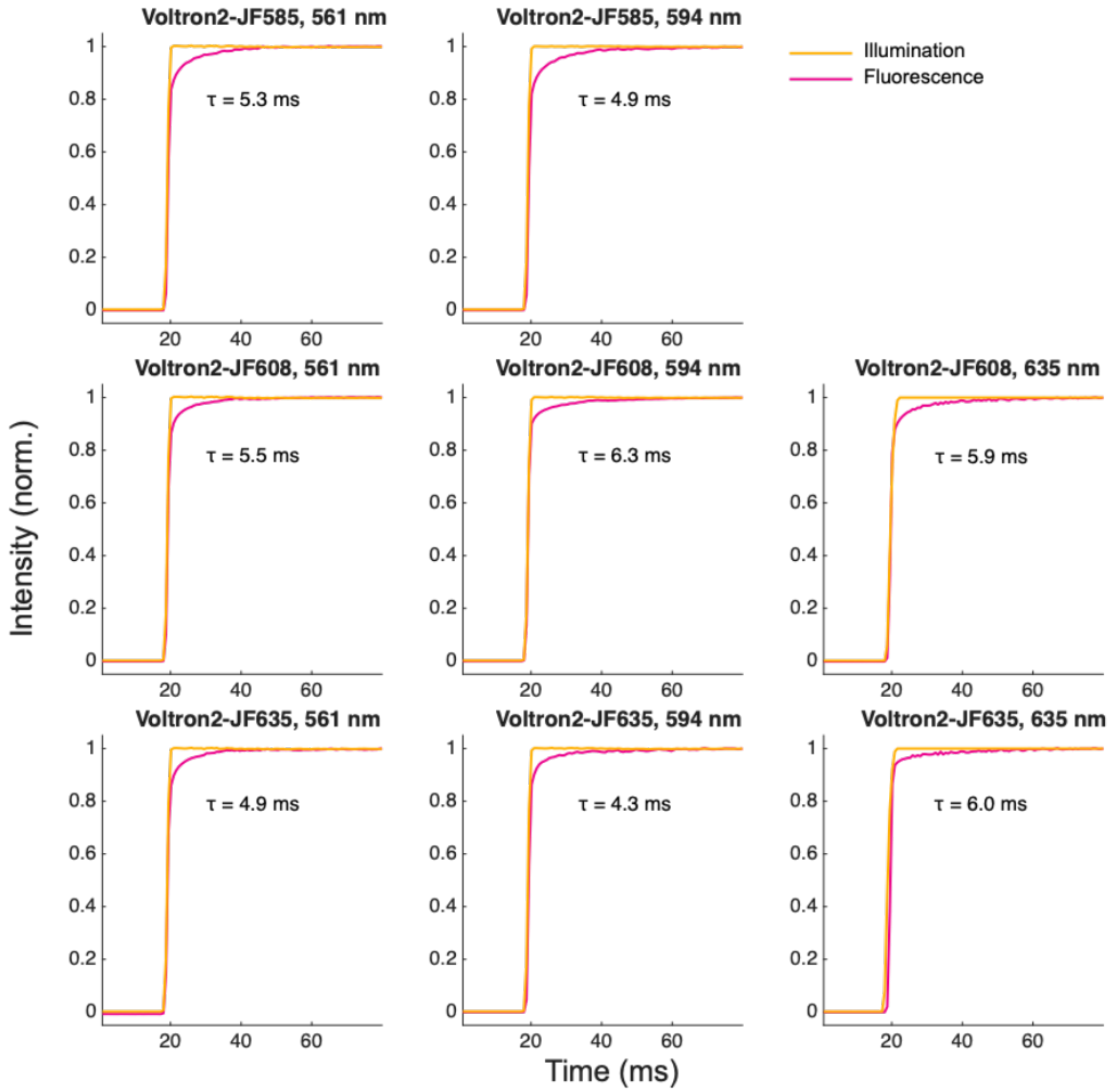


**Figure S11: Illumination-triggered fluorescence increase is shared across multiple excitation wavelengths and fluorophores.** HEK cells expressing Voltron2 labeled with HaloTag ligands JF585, JF608, or JF635 were tested under step illumination using different excitation wavelengths (561 nm at 150 mW/mm², 594 nm at 150 mW/mm², and 635 nm at 110 mW/mm²). The combination of JF585 with 635 nm excitation was omitted because this arrangement did not produce detectable fluorescence. In all conditions, fluorescence exhibited a characteristic millisecond-timescale upward relaxation upon illumination onset, indicating that Voltron2 shows photoactivation for multiple dyes and excitation wavelengths.

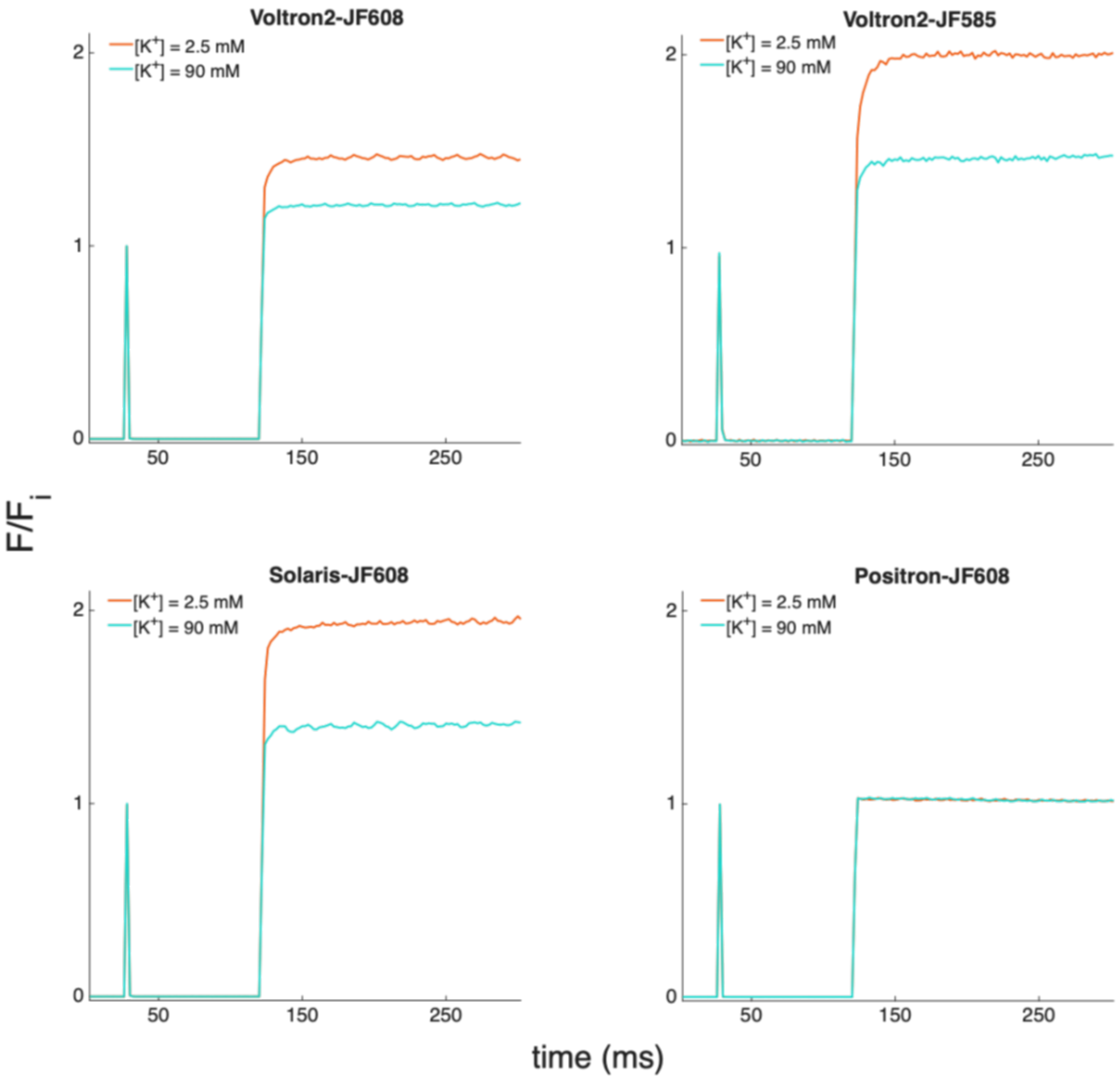


**Figure S12: Testing for VMAP mechanism across opsin-based GEVIs.** Hek cells expressed $K_{ir}2.1$ and the indicated FRET-opsin GEVI. The VMAP protocol was applied under extracellular $K^+$ concentrations of 2.5 mM and 90 mM, corresponding to membrane potentials of approximately -100 mV and -10 mV, respectively. Voltron2-JF608, Voltron2-JF585, and Solaris-JF608 each exhibited voltage-independent $F_i$ and an illumination-triggered activation into a voltage-sensitive state with fluorescence $F_v$. Positron-JF608 did not show a comparable photoactivation process that could serve as a voltage-insensitive reference (here fluorescence is normalized to $F_i = 1$, so the voltage-dependent change in Positron fluorescence does not appear on the plot).

| Figure | Excitation wavelength (nm) | Illumination intensity ($mW/mm^2$) | Frame rate (Hz) | Short pulse duration (ms) | Magnification (×) | Objective | Detector |
|---|---|---|---|---|---|---|---|
| 1B-G, S2, S4 | 607 | 100 | 1000 | 1 | 14 | Olympus XLPLN25XWMP2 25x, NA1.05 | Hamamatsu, ORCA-Fusion |
| 1I-L, 3B-C, 4, S9 | 607 | 100 | 166.7 | 1 | 14 | Olympus XLPLN25XWMP2 25x, NA1.05 | Hamamatsu, ORCA-Fusion |
| 2 3F-J, S7, S8 | 613 460 | 20 for 613 nm 2 for 460 nm | 166.7 | 0.5 | 2.3 | Olympus MVPLAPO 2XC, NA0.5 | Teledyne, Kinetix |
| S1 | 594 | 160 | 1000 | 0.2-1 | 20 | Olympus XLPLN25XWMP2 25x, NA1.05 | Hamamatsu, ORCA-Fusion |
| S5 | 594 | 160 for short pulse, 80 for long pulse | 1000 | 0.5 | 20 | Olympus XLPLN25XWMP2 25x, NA1.05 | Hamamatsu, ORCA-Fusion |
| S6 | 488, 594 | 3 for 488 nm 100 for 594 nm | 1000 | 0.5 | 20 | Olympus XLPLN25XWMP2 25x, NA1.05 | Hamamatsu, ORCA-Fusion |
| S11 | 561, 594, 635 | 150 for 561 nm, 150 for 594 nm, 110 for 635 nm | 1400 | N.A. | 6 | Olympus UIS2 UPlanSApo x20/0.75 | Hamamatsu, ORCA-Fusion |
| S12 | 561, 594 | 110 for 594 nm 40 for 561 nm | 1000 | 0.5 | 14 | Nikon N16XLWD-PF x 16/0.8 | Hamamatsu, ORCA-Fusion |

**Table S1: Imaging parameters for each figure panel.**

**Video S1: Optopatch VMAP video and representative traces of neuron culture without ML213 treatment.**

**Video S2: Optopatch VMAP video and representative traces of neuron culture with ML213 treatment.**

**Video S3: Timelapse VMAP video of hiPSC colonies imaged over 3 days.**

**Video S4: 3D VMAP image of a whole zebrafish embryo at 22 – 23 hpf.**

6. J. Zhang, D. Gong, A. Barrios, D. G. Itkis, T. Uenaka, M. Werning, R. Nehme, E. Fraenkel, S. Farhi, A. E. Cohen, A high-throughtput all-optical plateform for functional genomics of electrophysiology. *Manuscript in preparation*.

7. R. Tian, M. A. Gachechiladze, C. H. Ludwig, M. T. Laurie, J. Y. Hong, D. Nathaniel, A. V. Prabhu, M. S. Fernandopulle, R. Patel, M. Abshari, M. E. Ward, M. Kampmann, CRISPR

Interference-Based Platform for Multimodal Genetic Screens in Human iPSC-Derived Neurons. *Neuron* **104**, 239-255.e12 (2019).
